# Subfunctionalisation of paralogous genes and evolution of differential codon usage preferences: the showcase of polypyrimidine tract binding proteins

**DOI:** 10.1101/2020.08.30.274191

**Authors:** Jérôme Bourret, Fanni Borvető, Ignacio G. Bravo

## Abstract

Gene paralogs are copies of an ancestral gene that appear after gene or full genome duplication. When two sister gene copies are maintained in the genome, redundancy may release certain evolutionary pressures, allowing one of them to access novel functions. Here, we focused our study on gene paralogs on the evolutionary history of the three polypyrimidine tract binding protein genes (*PTBP*) and their concurrent evolution of differential codon usage preferences (CUPrefs) in vertebrate species.

*PTBP1-3* show high identity at the amino acid level (up to 80%), but display strongly different nucleotide composition, divergent CUPrefs and, in humans, distinct tissue-specific expression levels. Our phylogenetic inference results show that the duplication events leading to the three extant *PTBP1-3* lineages predate the basal diversification within vertebrates, and genomic context analysis illustrates that synteny has been well preserved over time for the three paralogs. We identify a distinct evolutionary pattern towards GC3-enriching substitutions in *PTBP1*, concurrent with an enrichment in frequently used codons and with a tissue-wide expression. In contrast, *PTBP2*s are enriched in AT-ending, rare codons, and display tissue-restricted expression. As a result of this substitution trend, CUPrefs are sharply different between mammalian *PTBP1*s and the rest of *PTBP*s. Genomic context analysis shows that GC3-rich nucleotide composition in *PTBP1*s is driven by local substitution processes, while the evidence in this direction is thinner for *PTBP2-3*. An actual lack of co-variation between the observed GC composition of *PTBP2-3* and that of the surrounding non-coding genomic environment would raise an interrogation on the origin of CUPrefs, warrantying further research on a putative tissue-specific translational selection. Finally, we communicate an intriguing trend for the use of the UUG-Leu codon, which matches the trends of AT-ending codons.

We interpret that our results are compatible with an scenario in which a combination of directional mutation–selection processes would have differentially shaped CUPrefs of *PTBPs* in Vertebrates: the observed GC-enrichment of *PTBP1* in Mammals may be linked to genomic location and to the strong and broad tissue-expression, while AT-enrichment of *PTBP2* and *PTBP3* would be associated with rare CUPrefs and thus, possibly to specialized spatio-temporal expression. Our interpretation is coherent with a gene subfunctionalisation process by differential expression regulation associated to the evolution of specific CUPrefs.

**1 Significance Statement:** In vertebrates, *PTBP* paralogs display strong differences in gene composition, gene expression regulation, and their expression in cell culture depends on their codon usage preferences. We show that placental mammals *PTBP1* have become GC-rich because of local substitution pressures, resulting in an enrichment of frequently used codons and in a strong, tissue-wide expression. On the contrary, *PTBP2* in vertebrates are AT-rich, with a lower contribution of local substitution processes to their specific nucleotide composition, show high frequency of rare codons and in placental mammals display a restricted expression pattern contrasting to that of *PTBP1*. The systematic study of composition and expression patterns of gene paralogs can help understand the complex mutation-selection interplay that shape codon usage bias in multicellular organisms.

## 2 Introduction

During mRNA translation ribosomes assemble proteins by specific amino acid linear polymerisation guided by the successive reading of mRNA nucleotide triplets, called codons. Each time a codon is read, it is chemically compared to the set of available tRNAs’ anticodons. Upon codon-anticodon match, the ribosome loads the tRNA and adds the associated amino acid to the nascent protein. The main 20 amino acids are encoded by 61 codons, so that multiple codons are associated with the same amino acid. These are named synonymous codons (Nirenberg and Matthaei, 1961; Khorana et al., 1966). Codon Usage Preferences (CUPrefs) refer to the differential usage of synonymous codons between species, between genes, or between genomic regions in the same genome (Grantham et al., 1980; Carbone et al., 2003). Mutation and selection are the two main forces shaping CUPrefs (Duret, 2002; Chamary et al., 2006; Plotkin and Kudla, 2011). Mutational biases relate to directional mechanistic biases during genome replication (Reijns et al., 2015; Apostolou-Karampelis et al., 2016), during genome repair (Lujan et al., 2012), or during recombination (Pouyet et al., 2017), preferentially introducing one nucleotide over others or inducing recombination and maintaining genomic regions depending on their composition. Mutational biases are well described in prokaryotes and eukaryotes, ranging from simple molecular preferences towards 3’A-ending in the Taq polymerase (Clark, 1988) to complex GC-biased gene conversion in vertebrates (Pouyet et al., 2017). Selective forces shaping CUPrefs are often described as translational selection. This notion refers to the ensemble of mechanistic steps and interactions during translation that are affected by the particular CUPrefs of the mRNA, so that the choice of certain codons at certain positions may actually enhance the translation process and can be subject to selection (Bulmer, 1991). Translational selection covers thus codon-independent effects on mRNA secondary structure, overall stability, and subcellular location (Presnyak et al., 2015; Novoa and Ribas de Pouplana, 2012), but also codon-mediated effects acting on mRNA maturation, programmed frameshifts, translation speed and accuracy, or protein folding (Caliskan et al., 2015; Mordstein et al., 2020; Spencer and Barral, 2012). Translational selection has been demonstrated in prokaryotes and in some eukaryotes (Satapathy et al., 2016; Percudani et al., 1997; Duret and Mouchiroud, 1999; Whittle and Extavour, 2016), often in the context of tRNA availability (Ikemura, 1981). However, its very existence in Vertebrates remains highly debated (Pouyet et al., 2017; Galtier et al., 2018).

Homologous genes share a common origin either by speciation (orthology) or by duplication events (paralogy) (Sonnhammer and Koonin, 2002). Upon gene (or full genome) duplication, the new genome will contain two copies of the original gene, referred to as in-paralogs. After speciation, each daughter cell will inherit one couple of paralogs, *i*.*e*. one copy of each ortholog (Koonin, 2005). The emergence of paralogs upon duplication may release the evolutionary constraints on the individual genes. Evolution can thus potentially lead to function specialisation, such as evolving a particular substrate preferences, or engaging each paralog on specific enzyme activity preferences in the case of promiscuous enzymes (Copley, 2020). Gene duplication can also allow one paralog to explore broader sequence space and to evolve radically novel functions, while the remaining counterpart can continue to assure the original function.

The starting point for our research are the experimental observations by Robinson and coworkers reporting differential expression of the polypyrimidine tract binding protein (*PTBP*) human paralogs as a function of their nucleotide composition (Robinson et al., 2008). Vertebrates genomes encode for three in-paralogous versions of the *PTBP* genes, all of them fulfilling similar functions in the cell: they form a class of hnRNP RNA-Binding Proteins that are involved in the modulation of mRNAs alternative splicing (Pina et al., 2018). Within the same genome, the three paralogs display high amino-acid sequence similarity, around 70% in humans, and with similar overall values in vertebrates (Pina et al., 2018).

Despite the high resemblance at the protein level, the three *PTBP* paralogs sharply differ in nucleotide composition, CUPrefs, and supposedly in tissue expression pattern. In humans *PTBP1* is enriched in GC3-rich synonymous codons and is widely expressed in all tissues, while *PTBP2* and *PTBP3* are AT3-rich and display an enhanced expression in the brain and in hematopoietic cells respectively (Supplementary Material, Figure S1). Robinson and coworkers studied the expression in human cells in culture of all three human *PTBP* paralogous genes placed under the control of the same promoter. They showed that the GC-rich paralog *PTBP1* was more highly expressed than the AT-rich ones, and that the expression of the AT-rich paralog *PTBP2* could be enhanced by synonymous codons recoding towards the use of GC-rich codons (Robinson et al., 2008). Here we have built on the evolutionary foundations of this observation and extended the analyses of CUPrefs to *PTBP* paralogs in vertebrate genomes. Our results suggest that paralog-specific directional changes in CUPrefs in mammalian *PTBP* concurred with a process of subfunctionalisation by differential tissue pattern expression of the three paralogous genes.

## 3 Material and Methods

### Sequence retrieval

We assembled a dataset of DNA sequences from 47 mammalian and 27 non-mammalian Vertebrates, and 3 from protostomes. Using the BLAST function on the nucleotide database of NCBI (NCBI Resource Coordinators, 2018) taking each of the human *PTBP* paralogs as references we looked for genes already annotated as *PTBP* orthologs (see supplementary Table S2 for accession numbers). We could retrieve the corresponding three orthologs in all Vertebrate species screened, except for the European rabbit *Oryctolagus cuniculus*, lacking *PTBP1*, and from the rifleman bird *Acanthisitta chloris*, lacking *PTBP3*. The final vertebrate dataset contained 75 *PTBP1*, 76 *PTBP2* and 75 *PTBP3* sequences. As outgroups for the analysis, we retrieved the orthologous genes from three protostome genomes, which contained a single *PTBP* homolog per genome. Our final dataset was consistent with the descriptions available in ENSEMBL and ORTHOMAM for the *PTBP* orthologs (Yates et al., 2020; Scornavacca et al., 2019; Pina et al., 2018). From the original dataset, we identified a subset of nine mammalian and six non-mammalian vertebrates species with a good annotation of the *PTBP* chromosome context. For these 15 species we retrieved synteny and composition information on the *PTBP* flanking regions and introns (Supplementary Table S3). Because of annotation hazards, intronic and flanking regions information were missing for some *PTBP*s in the African elephant *Loxodonta africana*, Schlegel’s Japanese Gecko *Gekko japonicus*, and the whale shark *Rhincodon typus* assemblies. For the selected 15 species the values for codon adaptation index (CAI) (Sharp and Li, 1987) and codon usage similarity index (COUSIN) (Bourret et al., 2019) were calculated using the COUSIN server (available at https://cousin.ird.fr) (Supplementary Table S4).

### Codon Usage analysis

For each *PTBP* gene we calculated codon composition, GC, GC3 and CUPrefs analyses via the COUSIN tool (Bourret et al., 2019). For each *PTBP* gene we constructed a vector of 59 positions with the relative frequencies of all synonymous codons. We applied different approaches to reduce information dimension for the analysis of CUPrefs, on the 229 59-dimension vectors: I) a k-means clustering; ii) a hierarchical clustering; and iii) a principal component analysis (PCA). Statistical analyses were performed using the ape and ade4 R packages and JMP v14.3.0. Correlation between matrices was assessed via the Mantel test. Non-parametric comparisons were performed using the Wilcoxon-Mann-Whitney test for assessing differences between the median values of the corresponding variable (either GC or GC3) among paralogs, and the Wilcoxon signed rank test for paired comparisons of the values for corresponding variable (either GC or GC3) for paralogs within the same genome. For the 15 species with well-annotated genomes we analyzed by a stepwise linear fit the correlation of paralog GC3 with two local compositional variables of the corresponding gene (GC content of intronic and flanking regions) and with three global compositional variables for the corresponding genomes (global GC3 in the complete genomic ORFome, global GC content in all introns, and global GC content in all flanking regions).

### Alignment and phylogenetic analyses

First, all sequences were aligned together, and we constructed a phylogenetic tree to verify whether each paralog assembly was monophyletic (Supplementary Figure S13). This was actually the case, and in this unbiased preliminary analysis all *PTBP1-3* were respectively monophyletic. Thus, to generate more robust alignments without introducing artefacts due to large evolutionary distances between in-paralogs, we proceeded stepwise, as follows: i) we aligned separately at the amino acid level each set of *PTBP* paralog sequences of mammals and non-mammalians Vertebrates; ii) for each *PTBP* paralog we merged the alignments for mammals and for non mammals, obtaining the three *PTBP1, PTBP2* and *PTBP3* alignments for all Vertebrates; iii) we combined the three alignments for each paralog into a single one; iv) we aligned the outgroup sequences to the global Vertebrate *PTBP*s alignment. All alignment steps were performed using MAFFT (Katoh et al., 2002). The final amino acid alignment was used to obtain the codon-based nucleotide alignment. The codon-based alignment was trimmed using Gblocks (Castresana, 2000) (Data available on Zenodo) Phylogenetic inference was performed at the amino acid and at the nucleotide level using RAxML v8.2.9, bootstrapping over 1000 cycles (Stamatakis, 2014). For nucleotides we used codon-based partitions and applied the GTR+G4 model while for amino acids we applied the LG+G4 model. For the 79 species used in the analyses we retrieved a species-tree from the TimeTree tool (Kumar et al., 2017). Distances between phylogenetic trees were computed using the Robinson-Foulds index, which accounts for differences in topology (Robinson and Foulds, 1981), and the K-tree score, which accounts for differences in both topology and branch length (Soria-Carrasco et al., 2007). We then calculated pairwise distances between branches on the nucleotide and amino acid based trees and compared them against CUPrefs-based pairwise distances to measure the impact of CUPrefs on the phylogeny. After phylogenetic inference, we computed marginal ancestral states for the respectively most recent common ancestors at the nucleotide level of each paralog, using RAxML. For each position the base with the maximum probability was used, and the sites for which RAxML could not infer with certainty the base were marked as missing data. We found 14%, 18% and 10% of missing bases respectively in *PTBP1, PTBP2* and *PTBP3*. Using these ancestral sequences we estimated the number of synonymous and non-synonymous substitutions of each extant sequence to the corresponding most recent common ancestor. We then compared the substitution matrices via a PCA analysis.

## 4 Results

### Vertebrate PTBP paralogs differ in nucleotide composition

In order to understand the evolutionary history of *PTBP* genes, we performed first a nucleotide composition and CUPrefs analysis on the three paralogs in 79 species. Overall, *PTBP1* are GC-richer than *PTBP2* and *PTBP3* (respective mean percentages 55.9, 42.3 and 44.9 for GC content and 69.5, 33.4 and 38.3 for GC3 content; Figure 1). In addition, *PTBP1*s show a difference in GC3 between mammalian and non-mammalian genes (respectively 79.8 against 59.9 mean percentages). A linear regression model followed by a Tukey’s honest significant differences analysis for GC3 using as explanatory levels paralog (*i*.*e. PTBP1-3*), taxonomy (*i*.*e*. mammalian or non-mammalian), and their interaction identifies three main groups of *PTBP*s (Table 1): a first one corresponding to mammalian *PTBP1*, a second one grouping non-mammalian *PTBP1*, and a third one encompassing all *PTBP2* and *PTBP3*. The largest explanatory factor for GC3 was the paralog *PTBP1-3*, accounting alone for 65% of the variance, while the interaction between the levels taxonomy and paralog captured around 15% of the remaining variance (Table 1). These trends are confirmed when performing paired comparisons between paralogs present in the same mammalian genome, with significant differences in GC3 content in the following order: *PTBP1* > *PTBP3* > *PTBP2* (Wilcoxon signed rank test: *PTBP1* vs *PTBP2*, mean diff=48.0, S=539.50, p-value <0.0001; *PTBP1* vs *PTBP3*, mean diff=43.5, S=517.50, p-value <0.0001; *PTBP3* vs *PTBP2*, mean diff=4.5, S=406.50, p-value <0.0001). Note that even if all of them significantly different, the mean paired differences in GC3 between *PTBP1* and *PTBP2-3* are ten times larger than the corresponding mean paired differences between *PTBP2* and *PTBP3*.

**Table 1:** Global linear regression model and post-hoc Tukey’s honest significant differences test for GC3 composition as explained variable and the explanatory levels paralog (*PTBP1-3*), taxonomy (*i*.*e*. mammalian or non-mammalian) and their interactions. Within each level, strata labelled with the same letter are not different from one another. Overall goodness of the fit: Adj Rsquare=0.83; F ratio=205.7; Prob > F: <0.0001.Individual effects for the levels: i) paralog: F ratio=274.3; Prob > F: <0.0001; ii) taxonomy: F ratio=27.2; Prob > F: <0.0001; iii) interaction paralog*taxonomy: F ratio=87.9; Prob > F: <0.0001.

| Level | Least Sq. Mean (GC3%) | Standard error | Tukey's HSD group |
| --- | --- | --- | --- |
| **Paralog** |  |  |  |
| *PTBP1* | 65.87 | 1.00 | A |
| *PTBP3* | 39.00 | 1.01 | B |
| *PTBP2* | 34.03 | 1.00 | C |
| **Taxonomy** |  |  |  |
| mammalian | 49.32 | 0.70 | A |
| non-mammalian | 43.28 | 0.92 | B |
| **Paralog\*Taxonomy** |  |  |  |
| *PTBP1*, mammalian | 79.81 | 1.22 | A |
| *PTBP1*, non-mammalian | 51.93 | 1.59 | B |
| *PTBP3*, non-mammalian | 41.64 | 1.62 | C |
| *PTBP3*, mammalian | 36.36 | 1.22 | C, D |
| *PTBP2*, non-mammalian | 36.27 | 1.59 | C, D |
| *PTBP2*, mammalian | 31.79 | 1.20 | D |

**Figure 1:**
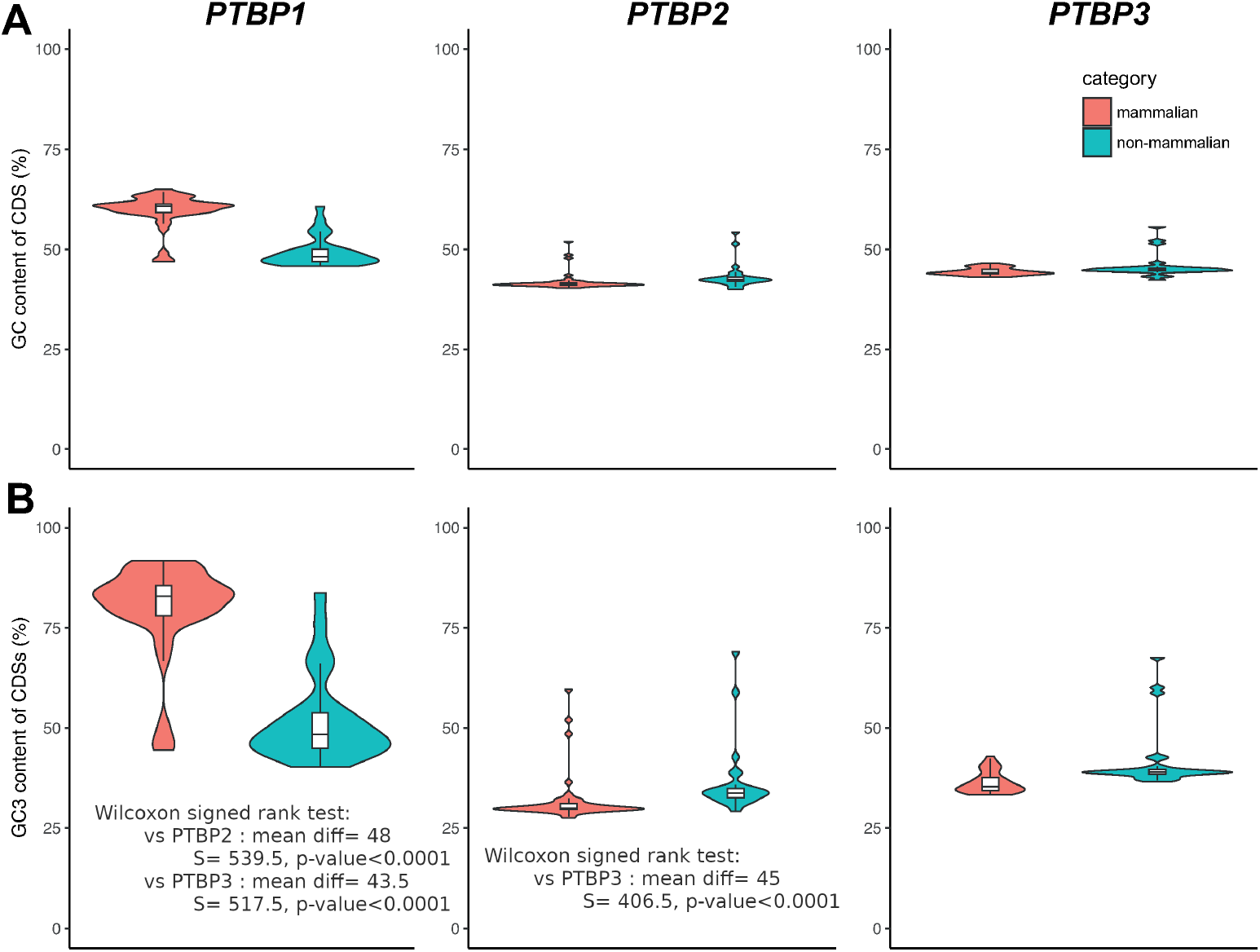
GC content (A) and GC3 content (B) of Vertebrates *PTBP*s. Violin plots display the overall distribution, while box and whiskers display median, quartiles and 95% of the corresponding values for mammalian (red) and non-mammalian (blue) individual genomes. The results of a the paired Wilcoxon signed rank tests between overall GC3 content of paralogs in the same genome are indicated in the inboxes.

After our model fit, an analysis of the distribution of the residuals between observed and expected values to the data allows to identify a number of outliers species with interesting taxonomical patterns in compositional deviation (Table 2). For non mammals, the three *PTBP* paralogs in the rainbow trout *Oncorhynchus mykiss* genome display high GC3 content (between 67% and 76%), all of them significantly higher than model-predicted values (expected values between 36% and 51%). A similar case occurs for the zebrafish *Danio rerio* genome: the three paralogs display GC3 values around 58%, which for *PTBP2* and *PTBP3* paralogs are significantly higher than predicted by the model (expected values around 38%). Very interestingly, for the monotrema platypus *Ornithorhynchus anatinus* as well as for the three marsupials in the dataset: the Tasmanian devil *Sarcophilus harrisii*, the koala *Phascolarctos cinereus* and the grey short-tailed opossum *Monodelphis domestica*, their *PTBP1* genes present similar GC3 content around 47%, which is significantly lower than predicted by the model (expected values around 79%).

**Table 2:** Individual genes with outlier values with respect to the linear regression expected values for the levels paralog (*PTBP1-3*), taxonomy (mammalian or non-mammalian) and their interactions.

| Species | paralog | observed GC3 (%) | expected GC3 (%) | deviation GC3 (%) |
| --- | --- | --- | --- | --- |
| **mammalian** |  |  |  |  |
| *Desmodus rotundus* | *PTBP2* | 59.60 | 31.79 | 27.81 |
| *Miniopterus natalensis* | *PTBP2* | 48.52 | 31.79 | 16.72 |
| *Monodelphis domestica* | *PTBP1* | 44.49 | 79.81 | -35.32 |
| *Ornithorhynchus anatinus* | *PTBP1* | 51.14 | 79.81 | -28.67 |
| *Ornithorhynchus anatinus* | *PTBP2* | 52.00 | 31.79 | 20.21 |
| *Phascolarctos cinereus* | *PTBP1* | 47.53 | 79.81 | -32.28 |
| *Sarcophilus harrisii* | *PTBP1* | 45.44 | 79.81 | -34.37 |
| **non-mammalian** |  |  |  |  |
| *Danio rerio* | *PTBP2* | 58.89 | 36.27 | 22.62 |
| *Danio rerio* | *PTBP3* | 60.08 | 41.64 | 18.44 |
| *Lepisosteus oculatus* | *PTBP3* | 58.73 | 41.64 | 17.10 |
| *Oncorhynchus mykiss* | *PTBP1* | 76.27 | 51.93 | 24.34 |
| *Oncorhynchus mykiss* | *PTBP2* | 69.03 | 36.27 | 32.76 |
| *Oncorhynchus mykiss* | *PTBP3* | 67.58 | 41.64 | 25.95 |
| *Pogona vitticeps* | *PTBP1* | 83.68 | 51.93 | 31.75 |

In many vertebrate species, strong compositional heterogeneities are observed along chromosomes with an arrangement of AT-rich and GC-rich regions, often referred to as “isochores”. To explore the influence of this genomic environment on the nucleotide composition of *PTBP*s, we analyzed for 15 species with well-annotated genomes the correlation of paralog GC3 with two local compositional variables of the corresponding gene (GC content of intronic and flanking regions) and with three global compositional variables for the corresponding genomes (global GC3 in the complete genomic ORFome, global GC content in all introns, and global GC content in all flanking regions)(Table 3 and Figure 2). First, for *D. rerio* the GC3 composition of *PTBP2* and *PTBP3* is clearly different from the rest, in line with the outlier results presented in Table 2. We have thus excluded the zebra fish values and performed an individual as well as a stepwise linear fit to explain the variance in GC3 composition by the variance in the local and global compositional variables mentioned above (Table 3). For all three *PTBP*s the local GC content explains best the corresponding GC3 content, but with strong differences between paralogs: while variation in the local composition captures almost perfectly variation in the GC3 content of *PTBP1* (R^2^=0.97) and relatively well in the case of *PTBP2* (R^2^=0.46), the fraction of variance explained by the local composition significantly drops for *PTBP3* (R^2^=0.15). It must be noted nevertheless that the GC3 variable ranges are different among paralogs, so that variation in GC3 values for *PTBP1* (roughly between 40% and 90%) is larger than for *PTBP2-3* (respectively 29%-38% and 34%-46%). This larger variable span in the case of *PTBP1* may allow for an increased power for detecting a significant correlation in composition values for this paralog.

**Table 3:** Results for an individual (left) or for a sequential (right) least squares regression for explaining variation in GC3 composition of *PTBP*s genes, by variation of different compositional variables, either local (introns or flanking regions of the corresponding gene) or global (all coding CDS, all introns and all flanking regions in the corresponding genome), in 14 well-annotated vertebrate genomes. For the sequential fit, variables are ordered according to their contribution to the sequentially better model for the corresponding paralog, and the order may thus differ between paralogs. Variables labelled with “n.s.” (not significant) do not contribute with significant additional explanatory power when added to the sequential model. BIC, Bayesian information content.

| *PTBP1* |  |  |  |  |  |
| --- | --- | --- | --- | --- | --- |
| Individual contributions |  |  | Sequential contribution |  |  |
| Parameter | R<sup>2</sup> | P value F test | Parameter | R<sup>2</sup> | BIC |
| Local_GC_intron | 0.9726 | <0.001 | Local_GC_intron | 0.9726 | 66.4765 |
| Local_GC_flanking | 0.5345 | 0.0069 | Local_GC_flanking | 0.974 (n.s.) | 68.3142 |
| Global_GC3_exome | 0.7279 | 0.0004 | Global_GC3_exome | 0.9749 (n.s.) | 70.3842 |
| Global_GC_introns | 0.116 | 0.2786 | Global_GC_flanking | 0.9803(n.s.) | 69.9886 |
| Global_GC_flanking | 0.1041 | 0.3065 | Global_GC_introns | 0.9806(n.s.) | 72.2531 |
| *PTBP2* |  |  |  |  |  |
| Individual contributions |  |  | Sequential contribution |  |  |
| Parameter | R<sup>2</sup> | P value F test | Parameter | R<sup>2</sup> | BIC |
| Local_GC_intron | 0.3738 | 0.0264 | Local_GC_flanking | 0.4558 | 60.1257 |
| Local_GC_flanking | 0.4558 | 0.0113 | Global_GC_introns | 0.4895(n.s.) | 61.8583 |
| Global_GC3_exome | 0.0943 | 0.3075 | Global_GC3_exome | 0.4914(n.s.) | 64.3761 |
| Global_GC_introns | 0.0488 | 0.4684 | Global_GC_flanking | 0.4934(n.s.) | 66.8894 |
| Global_GC_flanking | 0.0287 | 0.5801 | Local_GC_intron | 0.4974(n.s.) | 69.35 |
| *PTBP3* |  |  |  |  |  |
| Individual contributions |  |  | Sequential contribution |  |  |
| Parameter | R<sup>2</sup> | P value F test | Parameter | R<sup>2</sup> | BIC |
| Local_GC_intron | 0.1554 | 0.1825 | Local_GC_intron | 0.1554 | 74.7338 |
| Local_GC_flanking | 0.0522 | 0.4528 | Local_GC_flanking | 0.2095(n.s.) | 76.4388 |
| Global_GC3_exome | 0.0504 | 0.461 | Global_GC_introns | 0.2718(n.s.) | 77.9368 |
| Global_GC_introns | 0.0002 | 0.9661 | Global_GC3_exome | 0.2938(n.s.) | 80.1032 |
| Global_GC_flanking | 0.0024 | 0.8744 | Global_GC_flanking | 0.2938(n.s.) | 82.667 |

**Figure 2:**
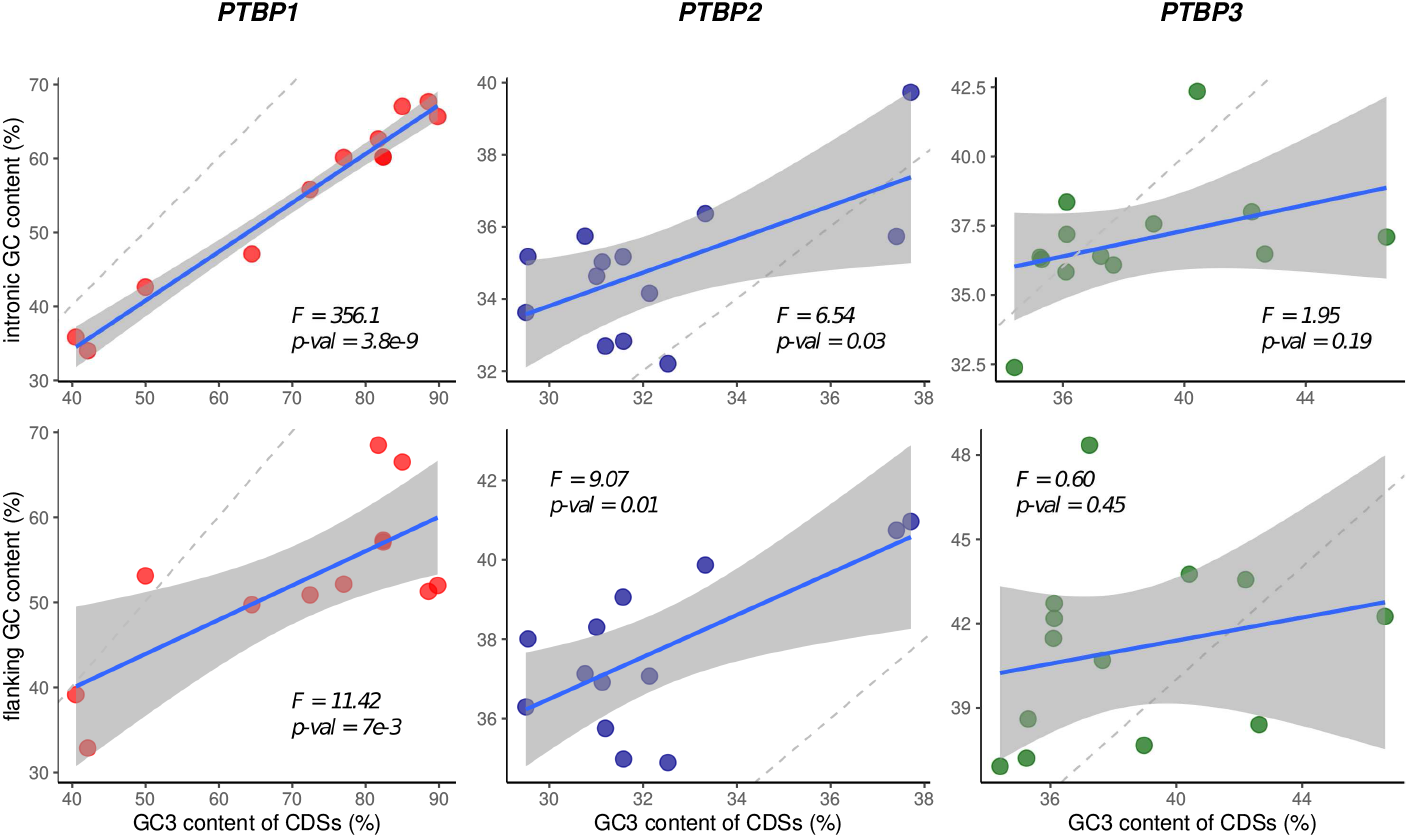
Variation in GC3 content of *PTBP*s (x-axis) and in the GC content of the corresponding introns (A, y axis) or flanking regions (B, y axis). Each dot represents one of the 15 individual genomes used for the genomic context analysis. For each graph, we performed a linear regression modelling (represented with the blue line for the fit and grey-shaded areas for the 95% confidence of the fit ; F-statistic and related p-values are given on the Figure); for each panel a grey line represents the *y* = *x* bisector.

### Vertebrate *PTBP* paralogs differ in CUPrefs

For each *PTBP* coding sequence we extracted the relative frequencies of synonymous codons and performed different approaches to reduce information dimension and visualise CUPrefs trends. The results of a principal component analysis (PCA) are shown in Figure 3 as well as in Supplementary Figure S5. The first PCA axis captured 68.9% of the variance, far before the second and the third axes (respectively 6.7% and 3.2%). Codons segregate in the first axis by their GC3 composition, the only exception being the UUG-Leu codon, which grouped together with AT-ending codons. This first axis differentiates mammalian *PTBP1*s on the one hand and *PTBP2*s and *PTBP3*s on the other hand. Non-mammalian *PTBP1*s scatter between mammalian *PTBP1*s and *PTBP3*s, along with the protostomates *PTBP*s. In the second PCA axis the only obvious (but nevertheless cryptic) codon-structure trends are: i) the split between C-ending and G-ending codons, but not between U-ending and A-ending codons; and ii) the large contribution in opposite directions to this second axis of the AGA and AGG-Arginine codons. This second PCA axis differentiates *PTBP2*s from *PTBP3*s paralogs, consistent with these composition trends. A paired-comparison confirms that *PTBP3*s are richer in C-ending codons than *PTBP2*s in the same genome, respectively 21.7% against 15.4% (Wilcoxon signed rank test: mean diff=6.2, S=1184.0, p-value <0.0001).

**Figure 3:**
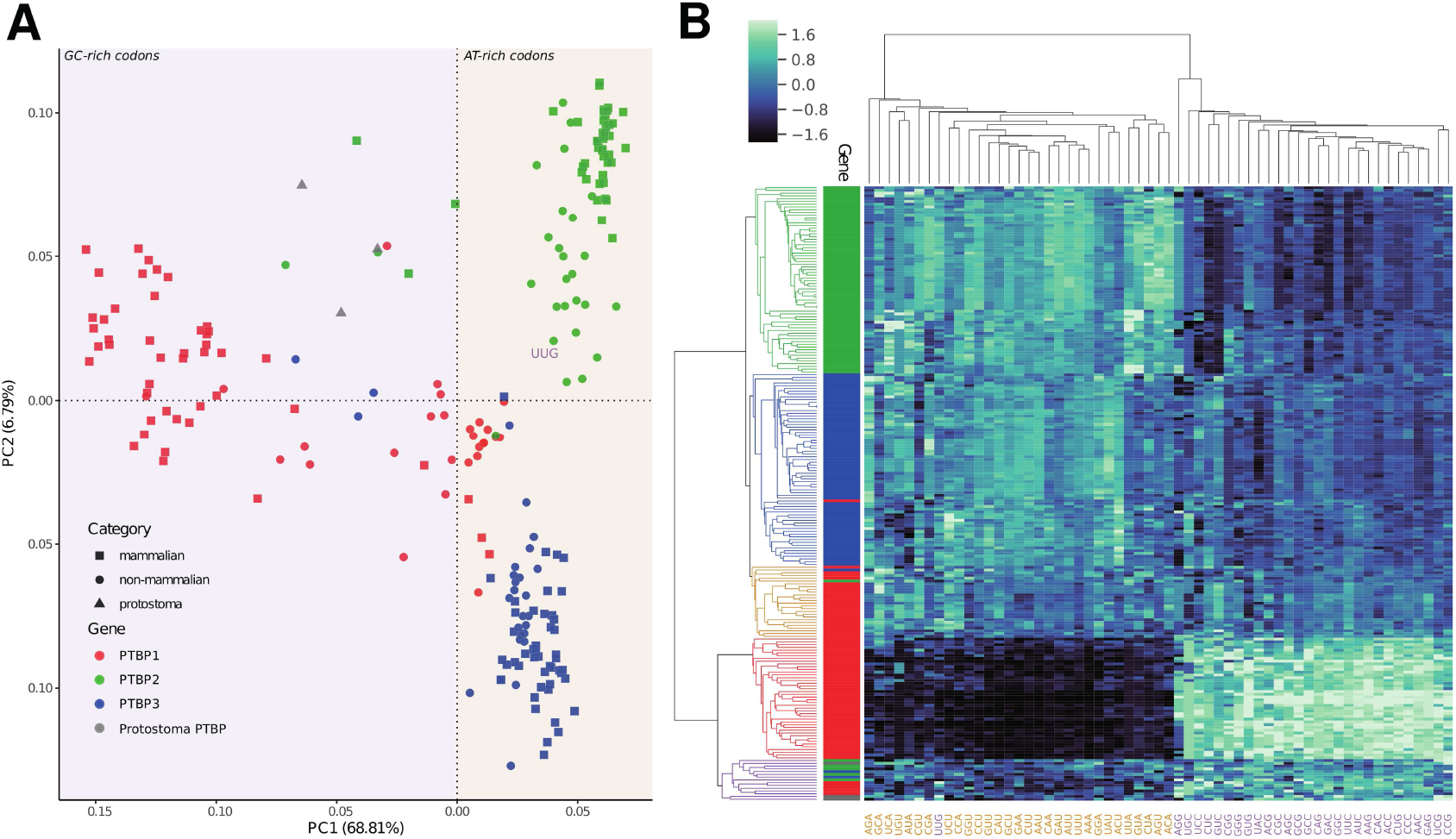
CUPrefs analysis of *PTBP*s. A) Plot of the two first dimensions of a PCA analysis based on the codon usage preferences of *PTBP1*s (red), *PTBP2*s (green), *PTBP3*s (blue) and protostoma (grey) individuals. Taxonomic information is included labellling mammals (squares), non-mammals (circles) and protostomes (triangles). The PCA was created using as variables the vectors of 59 positions (representing the relative frequencies of the 59 synonymous codons) for each individual gene. Shaded areas in purple (left) and orange (right) delimit the GC-rich and AT-rich grouping of codon variables according to the PCA. The UUG-Leu codon, colored in purple and placed on the Figure according to its eigenvalue, appears as a clear exception compared to the global trend of variables (Supplementary Figure S5) depicts a detailed positioning of the 59 PCA variables). The percentage of the total variance explained by each axis is shown in parenthesis. B) Heatmap of *PTBP*s individuals (rows) and synonymous codons (columns). Left dendrogram represents the hierarchical clustering of *PTBP*s based on their CUPrefs with colour codes that stand for the clusters created from this analysis. The side bar gives information on heatmap individuals regarding their origin : *PTBP1* (red), *PTBP2* (green), *PTBP3* (blue) or protostoma (grey). Note again the position of the UUG-Leu codon in the codon dendrogram, as the sole GC-ending codon clustering (in purple) with all other AT-ending codons (in orange)

As an additional way to identify groups of genes with similar CUPrefs, we applied a hierarchical clustering and a k-means clustering. Both analyses mainly aggregate *PTBP* genes by their GC3 richness. The *PTBP* dendrogram resulting of the hierarchical clustering shows five main clades that cluster the paralogs with a good match to the following groups: mammalian *PTBP1*s, non-mammalian *PTBP1*s, *PTBP2*s, *PTBP3*s and a fifth group containing the protostomata *PTBP*s and a few individuals of all three paralogs (rows in clustering in Figure 3; Kappa-Fleiss consistency score = 0.76). Regarding codon clustering, the hierarchical stratification sharply splits GC-ending codons from AT-ending codons, with the only exception again of the UUG-Leu codon, which consistently groups within the AT-ending codons. The elbow approach of k-means clustering identifies an optimal number of four clusters and separates the paralog genes with a good match as following: *PTBP1, PTBP2, PTBP3* and a group containing the protostomates and individuals from all paralogs (Kappa-Fleiss consistency score = 0.75).

Overall, k-means clustering and hierarchical clustering, both based on the 59-dimensions vectors of the CUPrefs, are congruent with one another (Kappa-Fleiss consistency score = 0.83), and largely concordant with the PCA results. CUPrefs define thus groups of *PTBP* genes consistent with their orthology and taxonomy. It is interesting to note that for some species the *PTBP* paralogs display unique distributions of CUPrefs, such as an overall similar CUPrefs in the three *PTBP* genes of the whale shark *Rhincodon typus*, or again some shifts in nucleotide composition between paralogs in the Natal long-fingered bat *Miniopterus natalensis*.

In order to characterise the directional CUPrefs bias of the different paralogs, we have analysed, for the 15 species with well-annotated genomes described above, the match between each individual *PTBP* and the average CUPrefs of the corresponding genome (Table 4). The COUSIN quantitative values compare the CUPrefs of a query sequence with those of a reference (in our case the coding genome of the corresponding organism), and can be directly interpreted in a qualitatively way, as described (Bourret et al., 2019). Briefly, COUSIN values around 1 reflect similar CUPrefs in the query sequence and in the reference, while values around 0 reflect CUPrefs close to random in the query sequence; COUSIN values above 1 reflect similar directional trends in CUPrefs in the query sequence and in the reference, but with stronger bias in the query sequence; COUSIN negative values reflect opposite CUPrefs between the query sequence and the reference. Our results highlight strong differences for mammalian paralogs: *PTBP1*s display COUSIN values above 1 while *PTBP2*s display COUSIN values below zero. The COUSIN results and interpretation are provided in (Supplementary Figure S14). These results mean that, in mammals, *PTBP1*s are enriched in commonly used codons in a higher proportion than the average in the genome, while *PTBP2*s are enriched in rare codons to the extent that their CUPrefs go in the opposite direction to the average in the genome. As for *PTBP3* in mammals, we observe COUSIN values below 0 in most cases or very close to 0 in the case of the horse *Equus caballus* and house mouse *Mus musculus*, implying a trend towards rare codons. In non-mammals however, *PTBP*s show an overall similarity to their respective reference genomic CUPrefs.

**Table 4:** Global linear regression model and post-hoc Tukey’s honest significant differences (HSD) test, the explained variable being the COUSIN value of the each *PTBP* gene compared with the average of the corresponding genome, and the explanatory levels paralog (*PTBP1-3*), taxonomy (*i*.*e*. mammalian or non-mammalian) and their interactions. Within each level, strata labelled with the same letter are not different from one another. Overall goodness of the fit: Adj Rsquare=0.82; F ratio=36.84; Prob > F: <0.0001.Individual effects for the levels: i) paralog: F ratio=40.72; Prob F: <0.0001; ii) taxonomy: F ratio=10.87; Prob > F: =0.0021; iii) interaction paralog*taxonomy: F ratio=28.11; Prob F: <0.0001.

| Level | Least Sq. Mean (COUSIN) | Standard error | Tukey's HSD group |
| --- | --- | --- | --- |
| **Paralog** |  |  |  |
| *PTBP1* | 1.45 | 0.11 | A |
| *PTBP3* | 0.29 | 0.11 | B |
| *PTBP2* | 0.19 | 0.11 | B |
| **Taxonomy** |  |  |  |
| mammalian | 0.44 | 0.080 | A |
| non-mammalian | 0.85 | 0.098 | B |
| **Paralog\*Taxonomy** |  |  |  |
| *PTBP1*, mammalian | 1.90 | 0.14 | A |
| *PTBP1*, non-mammalian | 0.99 | 0.17 | B |
| *PTBP2*, non-mammalian | 0.81 | 0.17 | B |
| *PTBP3*, non-mammalian | 0.75 | 0.17 | B |
| *PTBP3*, mammalian | -0.16 | 0.14 | C |
| *PTBP2*, mammalian | -0.43 | 0.14 | C |

### Phylogenetic reconstruction of *PTBP*s

We explored the evolutionary relationships between *PTBP*s by phylogenetic inference at the amino acid and at the nucleotide levels (Figure 4, Supplementary Figure S10). Our final dataset contained 74 *PTBP* sequences from mammals (47 species within 39 families) and non mammal vertebrates (27 species within 24 families). We used the *PTBP* genes from three protostome species as outgroup. Both amino acid and nucleotide phylogenies rendered three main clades grouping the *PTBP*s by orthology, so that all *PTBP1-3* orthologs were correspondingly monophyletic. In both topologies, *PTBP1* and *PTBP3* orthologs cluster together, although the protostome outgroups are linked to the tree by a very long branch, hampering the proper identification of the Vertebrate *PTBP* tree root. Amino acid and nucleotide subtrees were largely congruent (see topology and branch length comparisons in Table5). The apparently large nodal and split distance values between nucleotide and amino acid for *PTBP2* trees stem from disagreements in very short branches, as evidenced by the lowest K-tree score for this ortholog (as a reminder, the Robinson-Foulds index exclusively regards topology while the K-tree score combines topological and branch-length dependent distance between trees, see Material and Methods). In all three cases, internal structure of the ortholog trees essentially recapitulates species taxonomy at the higher levels (Table5). Some of the species identified by the regression analyses to display largely divergent nucleotide composition from the expected one given their taxonomy (Table 2) presented accordingly long branches in the phylogenetic reconstruction, such as *PTBP3* for *O. mykiss*, or rendered paraphyletic branching, as described above for *PTBP1* in marsupials and monotremes.

**Figure 4:**
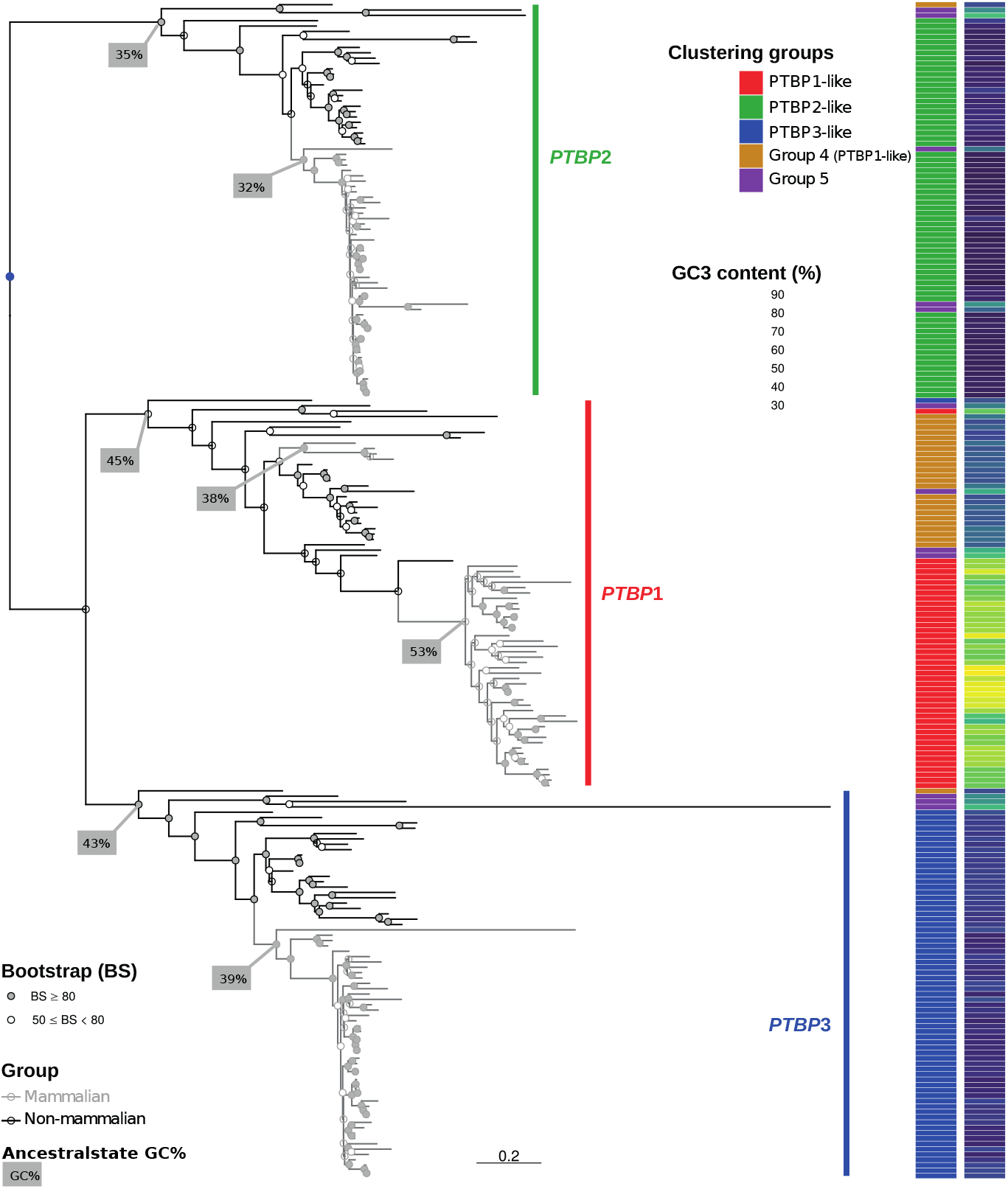
Maximum-likelihood nucleic acid phylogeny of *PTBP* genes. The phylogram depicts *PTBP2*s (green side bar), *PTBP1*s (red side bar) and *PTBP3*s (blue side bar) clades. The outgroup genes from protostomata are not shown to focus on the scale for vertebrate *PTBP*s, but their placement on the tree and the polarity they provide for vertebrate *PTBP*s is given by the blue dot. Gray branches indicate mammalian *PTBP*s, while black branches indicate non-mammalian species. Note the lack of monophyly for mammals for *PTBP1*s, with monotremes and marsupial lineages being paraphyletic to placental mammals. Filled dots on nodes indicate bootstrap values above 80, and empty dots indicate lower support values. Side bar on the left identifies the classification of each gene into the five groups identified by the hierarchical clusters, with the colour code in the inset. Side bar on the right displays GC3 content of the corresponding genes, with the gradient for the colour code ranging from 0 (blue) to 100% (yellow). The GC content of the main ancestral nodes is indicated in grey boxes.

**Table 5:** Comparison between species tree and the nucleotide based maximum likelihood tree for each *PTBP* paralog. The K-tree score compares topological and pairwise distances between trees after re-scaling overall tree length, with higher values corresponding to more divergent trees. The Robinson-Foulds score compares only topological distances between trees, the values shown correspond to the number of tree partitions that are not shared between two trees, so that higher values correspond to more divergent trees.

| Reference tree | Comparison tree | K-tree score | Robinson-Foulds score |
| --- | --- | --- | --- |
| **Nucleotide tree VS species tree** |  |  |  |
| *PTBP1* | Species tree | 0.759 | 42 |
| *PTBP2* | Species tree | 0.762 | 24 |
| *PTBP3* | Species tree | 1.700 | 28 |
| **Nucleotide tree VS Amino acid tree** |  |  |  |
| *PTBP1*-AA | *PTBP1*-NT | 0.149 | 78 |
| *PTBP2*-AA | *PTBP2*-NT | 0.129 | 110 |
| *PTBP3*-AA | *PTBP3*-NT | 0.380 | 40 |

We have then analysed the correspondence between nucleotide-based and amino acid-based pairwise distances to evaluate the impact of CUPrefs on the obtained phylogeny. We observe a good correlation between both reconstructions for all paralogs, except for mammalian *PTBP2*s, which display extremely low divergence at the amino acid level (see Figure 5 for values in mammalian paralogs, Supplementary Figure S8 for non-mammalian paralogs, and Supplementary Table S7 for the correlation between nucleotide-based and amino acid-based pairwise distances). For mammalian *PTBP1*s, the plot allows to clearly differentiate a cloud with the values corresponding to the monotremes+marsupial mammals, split apart from placental mammals in terms of both amino acid and nucleotide distances. This distribution matches well the fact that sequences from monotremes and marsupials cluster separately from placental mammals in the *PTBP1* phylogeny (see grey branches being paraphyletic for *PTBP1* in Figure 4). The same holds true for the platypus *PTBP3*, extremely divergent from the rest of the mammalian orthologs. The precise substitution patterns are analysed in detail below. The histograms describing the accumulation of synonymous and non-synonymous substitutions confirm that mammalian *PTBP1*s have accumulated the largest number of synonymous substitutions compared to non-mammalian *PTBP1*s and to other orthologs (Supplementary Figure S9).

**Figure 5:**
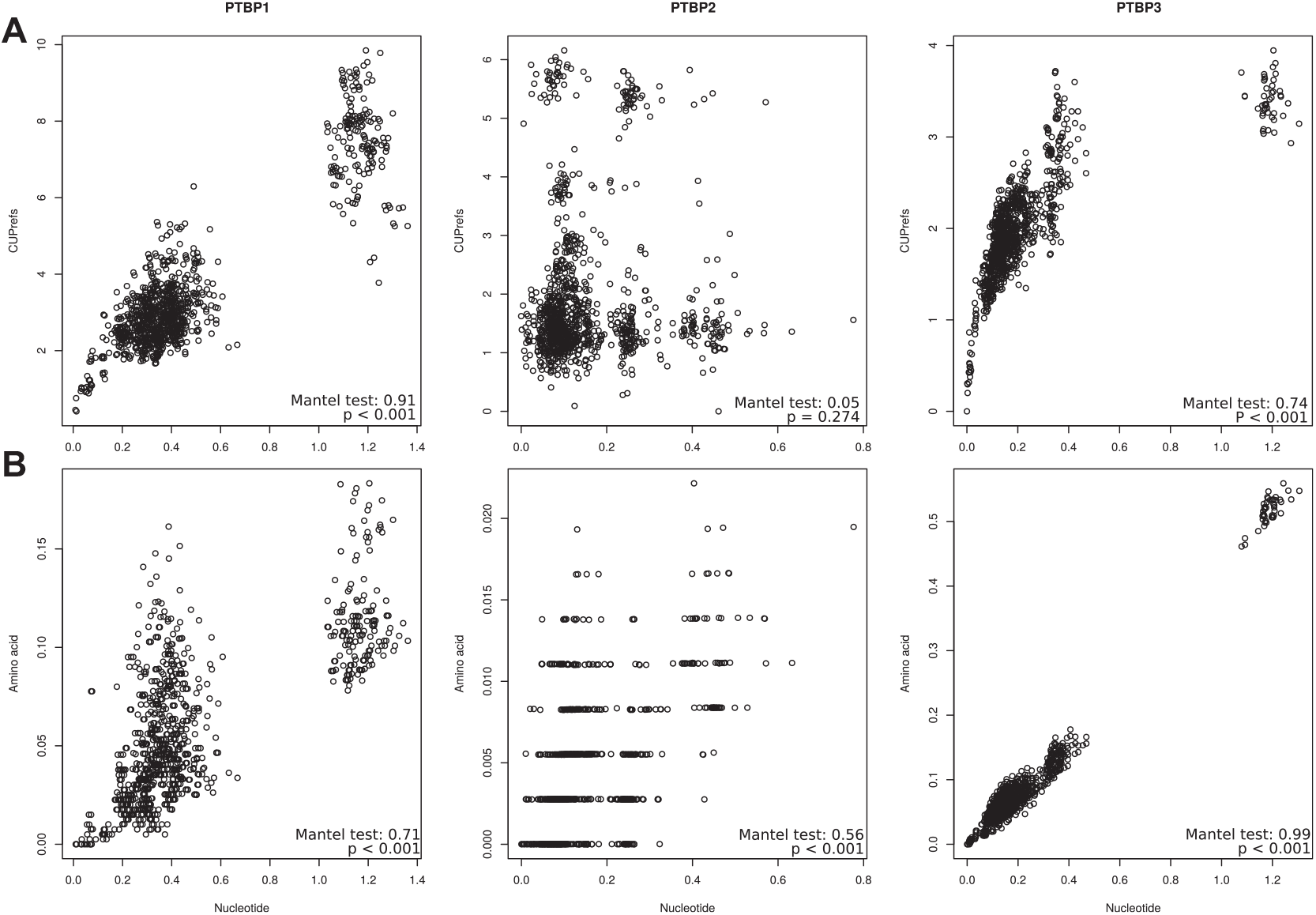
Nucleotide-based pairwise distances in the x-axis against A) CUPrefs-based and B) amino acid-based pairwise distances in the y-axis for the different mammalian *PTBP* orthologs. The results for a Mantel test assessing the correlation between the corresponding matrices are shown in each inset.

We have finally analysed the connection between nucleotide-based evolutionary distances within *PTBP* paralogs and CUPrefs-based distances (Figure 5 for mammalian paralogs and Supplementary Figure S8 for non-mammalian paralogs). A trend showing increased differences in CUPrefs as evolutionary distances increase is evident only for *PTBP1*s and *PTBP3*s in mammals. For mammalian *PTBP1*s the plot clearly differentiates a cloud with the values corresponding to monotremes and marsupials splitting apart from placental mammals in terms of both evolutionary distance and CUPrefs. For mammalian *PTBP2*s the plot captures the divergent CUPrefs of the platypus and of the bats *M. natalensis* and *Desmodus rotundus*, while for non-mammalian *PTBP2*s the divergent CUPrefs of the rainbow trout (*O. mykiss*) are obvious. Finally, for mammalian *PTBP3*s the large nucleotide divergence of the platypus paralog is evident. Importantly, all these instances of divergent behaviour (except for the platypus *PTBP3*) are consistent with the deviations described above from the expected composition by the mathematical modelling of the ortholog nucleotide composition (Table 2).

### Mammalian PTBP1s accumulate GC-enriching synonymous substitutions

We have shown that *PTBP1* genes are GC-richer and specifically GC3-richer than the *PTBP2* and *PTBP3* paralogs in the same genome, and that this enrichment is of a larger magnitude in placental *PTBP1*s. We have thus assessed whether a directional substitutional pattern underlies this enrichment, especially regarding synonymous substitutions. For this we have inferred the ancestral sequences of the respective most recent common ancestors of each *PTBP* paralog, recapitulated synonymous and non-synonymous substitutions between each *PTBP* individual and their ancestors, and constructed the corresponding substitution matrices (Table S11). The two first axes of a principal component analysis using these substitution matrices capture, with a similar share, 66.95% of the variance between individuals (Figure 6). The first axis of the PCA separates synonymous from non-synonymous substitutions. Intriguingly though, while T<->C transitions are associated with synonymous substitutions, as expected, G<->A transitions are instead associated with non-synonymous substitutions. The second axis separates substitutions by their effect on nucleotide composition: GC-stabilizing/enriching on one direction, AT-stabilizing/enriching on the other one. Strikingly, the substitutional spectrum of mammalian *PTBP1*s sharply differs from the rest of the paralogs. Substitutions in mammalian *PTBP1* towards GC-enriching changes, in both synonymous and non-synonymous compartments, are the main drivers of the second PCA axis. In contrast, synonymous substitutions in *PTBP3* as well as all substitutions in *PTBP2* tend to be AT-enriching. Finally, the substitution trends for *PTBP1* in mammals are radically different from those in non-mammals, while for *PTBP2* and *PTBP3*s the substitution patterns are similar in mammals and non-mammals for each of the compartments synonymous and non-synonymous.

**Figure 6:**
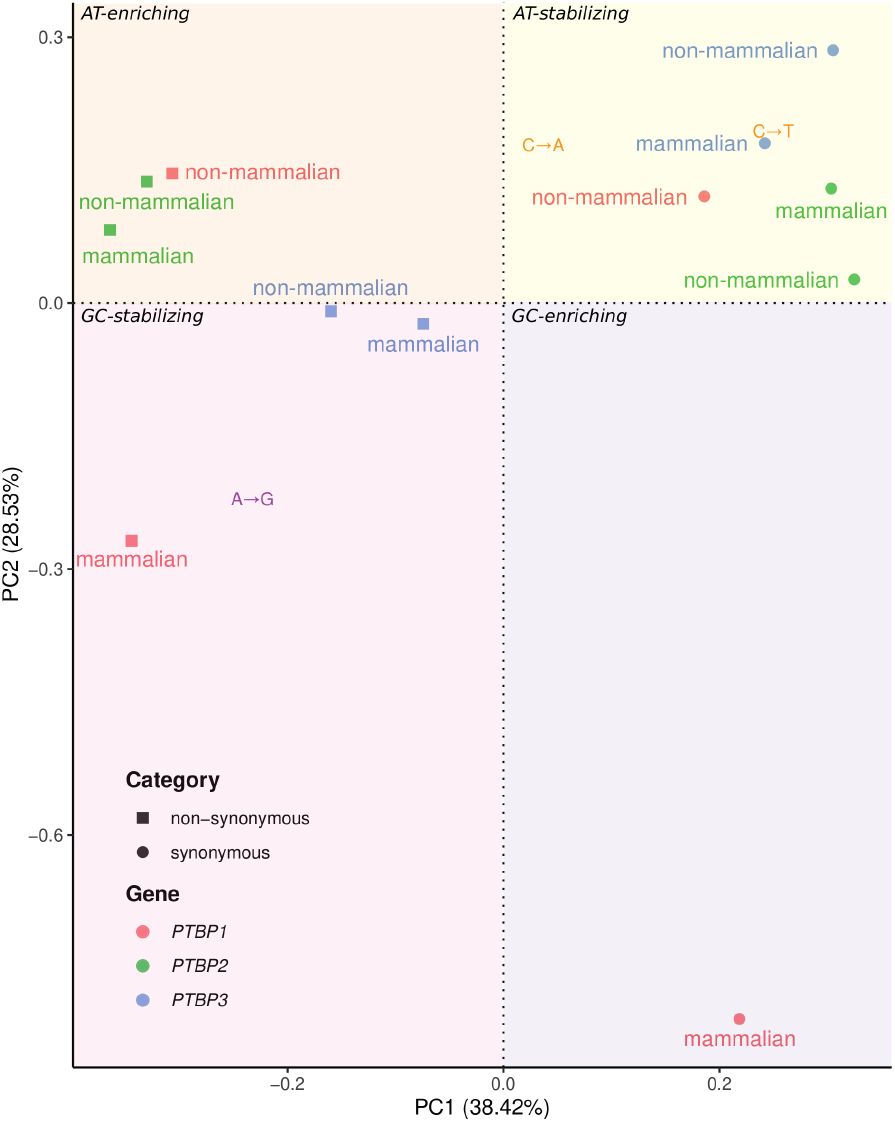
Spectra of synonymous and non-synonymous substitutions for *PTBP*s. This principal component analysis (PCA) has been built using the observed nucleotide synonymous and non-synonymous substitution matrices for each *PTBP* paralog, inferred after phylogenetic inference and comparison of extant and ancestral sequences. The variables in this PCA are the types of substitution (*e*.*g*. A->G), identified by a colour code as GC-enriching/stabilizing substitutions (purple and pink areas) or AT-enriching/stabilizing substitutions (orange and yellow areas). To facilitate the interpretation of the graph, all variables have been masked, except those that do not follow theses global, which have been plotted according to their eigenvalues (*i*.*e*. A->G, C->A and C->T) (all variables are shown unmasked in Supplementary Figure S15). Individuals in this PCA are the substitution categories in *PTBP* genes, stratified by their nature (synonymous or non-synonymous), by orthology (colour code for the different *PTBP*s is given in the inset) and by their taxonomy (mammals, or non-mammals).

## 5 Discussion

The non equal use of synonymous codons has puzzled biologists since it was first described. It has given rise to fruitful (and unfruitful) controversies between defenders of *all-is-neutralism* and defenders of *all-is-selectionism*, and has launched further the quest for additional molecular signaling beyond codons themselves (Callens et al., 2021). The main questions around CUPrefs are twofold. On the one hand, their origin: to what extent they are the result of fine interplay between mutation and selection processes. On the other hand, their functional implications: whether and how particular CUPrefs can be linked to specific gene expression regulation processes, broadly understood as downstream effects that modify the kinetics and dynamics of DNA transcription, mRNA maturation and stability, mRNA translation, and/or protein folding and stability. In the present work we have built on the experimental results of Robinson and coworkers, which communicated the differential expression of the *PTBP* human gene paralogs as a function of their CUPrefs (Robinson et al., 2008). From this particular example, we have aimed at exploring the nature of the connection between paralogous gene evolution and CUPrefs. Our results show that the three *PTBP* paralogous genes, which show divergent expression patterns in humans, also have divergent nucleotide composition and CUPrefs not just in humans but in most vertebrate species. We elaborate here on Robinson and coworkers experimental findings and propose here that this evolutionary pattern could be compatible with a phenomenon of phenotypic evolution by sub-functionalisation (in this case specialisation in tissue-specific expression levels), linked to genotypic evolution by association to specific CUPrefs patterns. Such conclusions invite to pursue Robinson and coworkers’ efforts by comparing *PTBP*s CUPrefs-modulated expression among numerous Vertebrate cell lines, especially between mammalians and non-mammalians ones. Consistent with studies on other paralog families (Munk et al., 2022), our results suggest, more generally, that a detailed analysis of differential CUPrefs in paralogs may help understand the divergent/convergent mutation-selection pressures that could underlie their functional differences.

We have reconstructed the phylogenetic relationships and analysed the evolution and diversity of CUPrefs among *PTBP* paralogs within 74 vertebrate species. The phylogenetic reconstruction shows that the genome of ancestral vertebrates already contained the three extant *PTBP* paralogs. This is consistent with the ortholog and paralog identification in the databases ENSEMBL and ORTHOMAM (Yates et al., 2020; Scornavacca et al., 2019; Pina et al., 2018). Although our results suggest that *PTBP1* and *PTBP3* are sister lineages, the distant relationship between the vertebrate genes and the protostome outgroup precludes the inference of a clear polarity between vertebrate *PTBP*s. We identify no instance of basal replacement between paralogs which may have appeared, for instance, as the replacement of an AT-rich paralog by a GC-rich one, leading to a loss of the AT-rich paralog and a duplication of the GC-rich one. Instead, the basal evolutionary histories of the different *PTBP*s comply well with those of the corresponding species. The most blatant mismatch between gene and species trees is the polyphyly of mammalian *PTBP1*s: monotremes and marsupials constitute a monophyletic clade, separate from placental mammals and not basal to them. Further, multiple findings in our results show sharp, contrasting patterns between *PTBP1* and the *PTBP2-3* paralogs: i) the excess of accumulation of synonymous substitutions in mammalian *PTBP1*s for a similar total number of changes (Supplementary Figure S9 and Table S11); ii) the larger differences in CUPrefs between genes with a similar total number of nucleotide changes in the case of *PTBP1*s in mammals (Figure 5 A); iii) the explicitly different spectrum of synonymous substitutions in *PTBP1*s, enriched in A->C, T->G and T->C changes (Figure 6); iv) the sharp difference of CUPrefs between *PTBP1*s and *PTBP2-3*s; and v) the clustering of *PTBP1* genes in monotremes and marsupials together with *PTBP1* genes in non-mammals according to their CUPrefs (Figure3 A). Overall, the particular nucleotide composition and the associated CUPrefs in mammalian *PTBP1* genes are most likely associated to specific local substitution biases as shown by the strong correlation between coding and non-coding GC content in *PTBP1* orthologs, while CUPrefs in *PTBP2-3*s cannot be explained alone by such local substitution biases (Figure 2; Table 3).

While GC3-rich nucleotide composition and CUPrefs of mammalian *PTBP1*s are dominated by local substitution biases, this is not the case for mammalian *PTBP2*, overall AT3-richer and without any clear correlation between coding and non-coding GC content among studied species (Figure 2; 3). As mentioned above, a note of caution should be raised here, as the variable range for GC composition among *PTBP1*s is larger than for *PTBP2-3*s, so that covariation analyses may have less power for the latter paralogs. In vertebrates, nucleotide composition varies strongly along chromosomes, so that long chromatin stretches, historically named “isochores”, appear enriched in GC or in AT nucleotides and present particular physico-chemical profiles (Caspersson et al., 1968). Local mutational biases and GC-biased gene conversion mechanism may underlie such heterogeneity, predominantly shaping local nucleotide composition in numerous Vertebrates genomes, so that the physical location of a gene along the chromosome largely explains its CUPrefs (Holmquist, 1989). In agreement with these hypotheses for local mutational biases, variation in GC3 composition of *PTBP1*s is almost totally (R2=0.97) explained by the variation in local GC composition (Figure 2; Table 3), suggesting that a similar substitution bias has shaped the GC-rich composition of the flanking, intronic and coding regions of *PTBP1*s. The same trend, albeit to a lesser degree holds also true for *PTBP2*s (R2=0.45). GC-biased gene conversion is often invoked as a powerful mechanism underlying such local GC-enrichment processes, leading to the systematic replacement of the alleles with the lowest GC composition by a GC-richer homolog (Marais, 2003). It has been proposed that gene expression during meiosis (evaluated as mRNA detection) correlates with a decreased probability of GC-biased gene conversion during meiotic recombination (Pouyet et al., 2017). Expression of *PTBP1* in human cells is documented during meiosis in the ovocite germinal line and expression of the AT-rich *PTBP2* has been observed during spermatogenic meiosis (Zagore et al., 2015; Hannigan et al., 2017). Expression during meiosis might thus have hindered GC-biased gene conversion for *PTBP1-2*s, provided that this expression pattern observed in humans was displayed also by the mammalian ancestor and that it is shared between mammalian species. With these assumptions, and thus, with caution, the GC-richness of *PTBP1* cannot be accounted for by GC-biased gene conversion, while the low GC content of *PTBP2* could be explained by an accumulation of GC->AT and AT->AT substitutions. All this notwithstanding, our results shot that GC3 enrichment in mammalian *PTBP1* and the concurrent trend for enriched use of common codons are associated mostly with placental mammals, and that non-placental mammals display divergent composition and differ from the model expectations. This synapomorphy of a sudden change in nucleotide composition is strongly compatible with a GC-biased gene conversion event in the placental ancestor that may have led to fixation of the ancestral version of the extant GC-rich *PTBP1*. Regarding *PTBP3*, the low GC-content together with the low correlation with either coding nor non-coding local GC-content could indicate that other mechanisms may shape the observed CUPrefs for this paralog.

In mammals, global GC-enriching genomic biases strongly impact CUPrefs, so that the most used codons in average tend to be GC-richer (Hershberg and Petrov, 2009). For this reason, mammalian GC3-rich *PTBP1*s match better the average genomic CUPrefs than AT3-richer *PTBP2* and *PTBP3*, which display CUPrefs in the opposite direction to the average of the genome. In the case of humans, *PTBP1* presents a COUSIN value of 1.75, consistent with a substantial enrichment in preferentially-used codons, while on the contrary, the COUSIN values of -0.48 for *PTBP2* and of -0.23 for *PTBP3* point towards a strong enrichment in rare codons (Supplementary Table S4). The poor match between human *PTBP2* CUPrefs and the human average CUPrefs could result in low expression of these genes in different human and murine cell lines, otherwise capable of expressing *PTBP1* at high levels and of expressing *PTBP3* at a lesser degree (Robinson et al., 2008). The barrier to *PTBP2* expression seems to be the translation process, as *PTBP2* codon-recoding towards GC3-richer codons results in strong protein production in the same cellular context, without significant changes in the corresponding mRNA levels (Robinson et al., 2008). Such codon recoding strategy towards preferred codons has become a standard practice for gene expression engineering that provides with very good expression results, despite our lack of understanding about the whole impact of local and global gene composition, nucleotide CUPrefs, and mRNA structure on gene expression (Brule and Grayhack, 2017).

The poor expression ability of *PTBP2* in human cells, the increase in protein production by the introduction of common codons, along with substitution biases failing to explain entirely *PTBP2* nucleotide composition and CUPrefs, raise the question of the adaptive value of poor CUPrefs in this paralog. Specific tissue-dependent or cell-cycle dependent gene expression regulation patterns have been invoked to explain the codon usage-limited gene expression for certain human genes, such as *TLR7* or *KRAS* (Newman et al., 2016; Lampson et al., 2013; Fu et al., 2018). The expression levels of the three *PTBP* paralogs are tissue-dependent in humans (Supplementary Figure S1), and through mammals (Keppetipola et al., 2012; Wagner and Garcia-Blanco, 2002; Spellman et al., 2007). In the case of the duplicated genes, subfunctionalisation through specialisation in spatio-temporal gene expression has been proposed as the main evolutionary force driving conservation of paralogous genes (Ferris and Whitt, 1979). Such differential gene expression regulation in paralogs has actually been documented for a number of genes at very different taxonomic levels (Donizetti et al., 2009; Guschanski et al., 2017; Freilich et al., 2006). Specialised expression patterns in time and space can result in antagonistic presence/absence of the paralogous proteins (Adams et al., 2003). This is precisely the case of *PTBP1* and *PTBP2* during human central nervous system development: in non-neuronal cells, *PTBP1* represses *PTBP2* expression by the skip of the exon 10 during *PTBP2* mRNA maturation, while during neuronal development, the micro RNA miR124 down-regulates *PTBP1* expression, which in turn leads to up-regulation of *PTBP2* (Keppetipola et al., 2012; Makeyev et al., 2007). Regarding non-human species, the available data about tissue-dependent and/or ontogeny-dependent differential expression at the transcription level (Abugessaisa et al., 2021) are largely concordant with the human data for *PTBP*, showing a tissue-wide transcription of *PTBP1*, a more restricted one for *PTBP3* together with an enrichment of *PTBP2* transcription in the central nervous system, as exemplified in the mouse (Barbosa-Morais et al., 2012), in the rat (Yu et al., 2014), in the cow (Merkin et al., 2012), in the gray short-tailed opossum (Brawand et al., 2011), or in the chicken (Barbosa-Morais et al., 2012). Finally, despite the high level of amino acid similarity between both proteins, *PTBP1* and *PTBP2* seem to perform complementary activities in the cell and to display different substrate specificity, so that they are not directly inter-exchangeable by exogenous manipulation of gene expression patterns (Vuong et al., 2016).

In addition to local genomic context analyses, we explored *PTBP* chromosomal location and local synteny (Figure 7). The results show that, while it is clear that the position of human *PTBP1* is telomeric and thus in one of the GC-richer region of human chromosome 9, most *PTBP*s do not map to the telomeres. Therefore, while the specific location of human *PTBP1* may have influenced its CUPrefs, it is unclear whether the chromosomic location of *PTBPs* have an impact on observed nucleotide composition. Synteny of *PTBP*s genes seems to be conserved, with some exceptions: most mammalian *PTBP1*s have a conserved synteny block that differs from non-mammalian species, with the exception of *D. rerio*. For *PTBP2* and *PTBP3* synteny seems conserved between mammalian and non-mammalian species again with the exception of *D. rerio*, lacking the *SUSD1* gene between *PTBP3* and *UGCG*. Such results could indicate that vertebrate radiation has been followed up by a change of *PTBP1* genomic context, with a swapping in flanking genes in mammalian branches. These results could be related to the observed *PTBP1* differential GC-content between mammalian and non-mammalian species.

**Figure 7:**
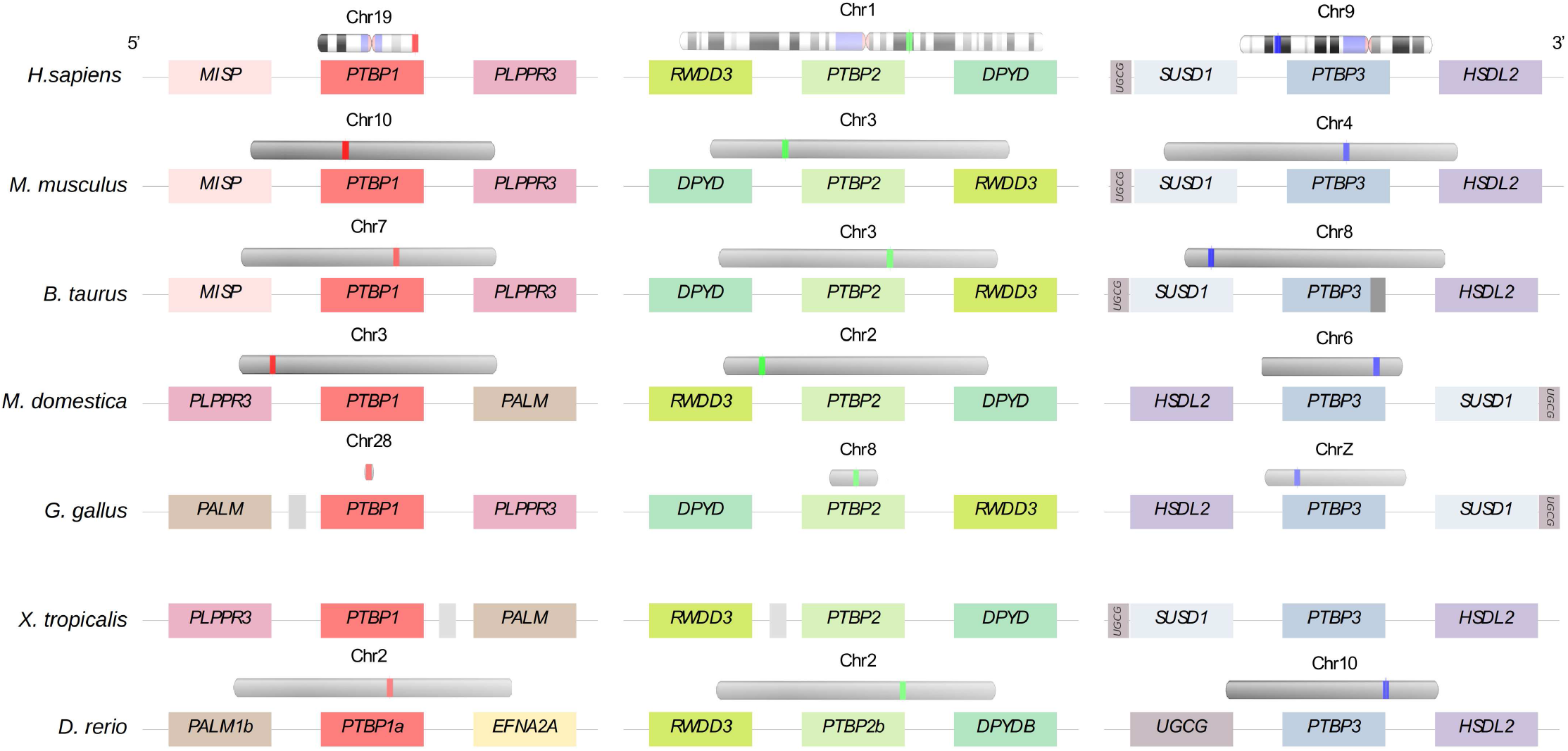
Placement on the chromosomes and genomic context of the three *PTBP* paralogs in a subset of the studied species.

In a different subject, we want to drive the attention of the reader towards the puzzling trend of the UUG-Leu codon in our CUPrefs analyses. This UUG codon is the only GC-ending codon systematically clustering with AT-ending codons in all our analyses, and not showing the expected symmetrical behaviour with respect to UUA-Leu (see Figure 3). Such behaviour for UUG has been depicted, but not discussed, in other analyses of CUPrefs in mammalian genes (see figure 7 in Laurin-Lemay et al. (2018)), in coronavirus genomes (Daron and Bravo, 2021), in plants (Clément et al., 2017) as well as for AGG-Arg and GGG-Gly in a global study of codon usages across the tree of life (see figure 1 in (Novoa et al., 2019)). The reasons underlying the clustering of UUG with AT-ending codons are unclear. A first line of thought could be functional: the UUG-Leu codon is particular because it can serve as alternative starting point for translation (Peabody, 1989). However, other codons such as ACG or GUG act more efficient than UUG as alternative translation initiation, and do not display any noticeable deviation in our results (Ivanov et al., 2011). A second line of thought could be related to the tRNA repertoire, but both UUG and UUA are decoded by similar numbers of dedicated tRNAs in the vast majority of genomes (*e*.*g*. respectively six and seven tRNA genes in humans (Palidwor et al., 2010)). Finally, another line of thought suggests that UUG and AGG could be disfavoured if substitution pressure towards GC is very high, despite being GC-ending codons (Palidwor et al., 2010). Indeed, the series of synonymous transitions UUA->UUG->CUG for Leucine and the substitution chain AGA->AGG->CGG for Arginine are expected to lead to a depletion of UUG and of AGG codons when increasing GC content. Both UUG and ACG codons would this way display a non-monotonic response to GC-substitution biases (Palidwor et al., 2010). In our data-set, however, AGG maps with the rest of GC-ending codons, symmetrically opposed to AGA as expected, and strongly contributing to the second PCA axis. Thus, only UUG displays frequency patterns similar to those of AT-ending codons. We humbly admit that we do not find a satisfactory explanation for this behaviour and invite researchers in the field to generate alternative explanatory hypotheses.

We have presented here an evolutionary analysis of the *PTBP* paralogs family as a showcase of CUPrefs evolution upon gene duplication. Our results show that differential nucleotide composition and CUPrefs in *PTBPs*s have evolved in parallel with differential gene expression regulation patterns. In the case of *PTBP1*, the most tissue-wise expressed of the paralogs, we have potentially identified compositional and substitution biases as the driving force leading to strong enrichment in GC-ending codons. In contrast, for *PTBP2* the enrichment in AT-ending codons is rather compatible with selective forces related to specific spatio-temporal gene expression pattern, antagonistic to those of *PTBP1*. Our results suggest that the systematic study of composition, genomic location and expression patterns of paralogous genes can contribute to understanding the complex mutation-selection interplay shaping CUPrefs in multicellular organisms.

## 6 Acknowledgments

J.B. was the recipient of a PhD fellowship from the French Ministry of Education and Research. This study was supported by the European Union’s Horizon 2020 research and innovation program under the grant agreement CODOVIREVOL (ERC-2014-CoG-647916) to I.G.B. The authors acknowledge the CNRS and the IRD for additional intramural support. The computational results presented have been achieved in part using the IRD Bioinformatic Cluster itrop.

## 7 Data Availability Statement

All data required to reproduce our findings is available on zenodo (https://doi.org/10.5281/zenodo.5789766), or provided in the tables in the main text and in the Supplementary Material section.

## 8 Disclosure

I.G.B. is a PCI recommender.

## Notes

https://doi.org/10.5281/zenodo.5789766

### Competing Interest Statement

The authors have declared no competing interest.

### Summary of Updates

-Phrasing to clarify our interpretations -updated material and methods to contain more details -updated phylogeny figure and its description -added a figure (Figure 7, previously in supplementary) -typos

## References

Abugessaisa I, Ramilowski JA, Lizio M, Severin J, Hasegawa A, Harshbarger J, Kondo A, Noguchi S, Yip CW, Ooi J, Tagami M, Hori F, Agrawal S, Hon C, Cardon M, Ikeda S, Ono H, Bono H, Kato M, Hashimoto K, Bonetti A, Kato M, Kobayashi N, Shin J, de Hoon M, Hayashizaki Y, Carninci P, Kawaji H, Kasukawa T. 2021, January. FANTOM enters 20th year: expansion of transcriptomic atlases and functional annotation of non-coding RNAs. Nucleic Acids Research. 49(D1):D892–D898.

Adams KL, Cronn R, Percifield R, Wendel JF. 2003, April. Genes duplicated by polyploidy show unequal contributions to the transcriptome and organ-specific reciprocal silencing. Proceedings of the National Academy of Sciences of the United States of America. 100(8):4649–4654.

Apostolou-Karampelis K, Nikolaou C, Almirantis Y. 2016, August. A novel skew analysis reveals substitution asymmetries linked to genetic code GC-biases and PolIII a-subunit isoforms. DNA research: an international journal for rapid publication of reports on genes and genomes. 23(4):353–363.

Barbosa-Morais NL, Irimia M, Pan Q, Xiong HY, Gueroussov S, Lee LJ, Slobodeniuc V, Kutter C, Watt S, Colak R, Kim T, Misquitta-Ali CM, Wilson MD, Kim PM, Odom DT, Frey BJ, Blencowe BJ. 2012, December. The evolutionary landscape of alternative splicing in vertebrate species. Science (New York, N.Y.). 338(6114):1587–1593.

Bourret J, Alizon S, Bravo IG. 2019, December. COUSIN (COdon Usage Similarity INdex): A Normalized Measure of Codon Usage Preferences. Genome Biology and Evolution. 11(12):3523–3528. Publisher: Oxford Academic.

Brawand D, Soumillon M, Necsulea A, Julien P, Csárdi G, Harrigan P, Weier M, Liechti A, Aximu-Petri A, Kircher M, Albert FW, Zeller U, Khaitovich P, Grützner F, Bergmann S, Nielsen R, Pääbo S, Kaessmann H. 2011, October. The evolution of gene expression levels in mammalian organs. Nature. 478(7369):343–348.

Brule CE, Grayhack EJ. 2017. Synonymous Codons: Choose Wisely for Expression. Trends in genetics: TIG. 33(4):283–297.

Bulmer M. 1991, November. The selection-mutation-drift theory of synonymous codon usage. Genetics. 129(3):897–907.

Caliskan N, Peske F, Rodnina MV. 2015, May. Changed in translation: mRNA recoding by 1 programmed ribosomal frameshifting. Trends in Biochemical Sciences. 40(5):265–274.

Callens M, Pradier L, Finnegan M, Rose C, Bedhomme S. 2021. Read between the lines: Diversity of nontranslational selection pressures on local codon usage. Genome Biology and Evolution. 13.

Carbone A, Zinovyev A, Képès F. 2003, November. Codon adaptation index as a measure of dominating codon bias. Bioinformatics (Oxford, England). 19(16):2005–2015.

Caspersson T, Farber S, Foley GE, Kudynowski J, Modest EJ, Simonsson E, Wagh U, Zech L. 1968, January. Chemical differentiation along metaphase chromosomes. Experimental Cell Research. 49(1):219–222.

Castresana J. 2000, April. Selection of conserved blocks from multiple alignments for their use in phylogenetic analysis. Molecular Biology and Evolution. 17(4):540–552.

Chamary JV, Parmley JL, Hurst LD. 2006, February. Hearing silence: non-neutral evolution at synonymous sites in mammals. Nature Reviews. Genetics. 7(2):98–108.

Clark JM. 1988, October. Novel non-templated nucleotide addition reactions catalyzed by procaryotic and eucaryotic DNA polymerases. Nucleic Acids Research. 16(20):9677–9686.

Clément Y, Sarah G, Holtz Y, Homa F, Pointet S, Contreras S, Nabholz B, Sabot F, Sauné L, Ardisson M, Bacilieri R, Besnard G, Berger C Angélique Cardi, De Bellis F, Fouet O, Jourda C, Khadari B, Lanaud C, Leroy T, Pot D, Sauvage C, Scarcelli N, Tregear J, Vigouroux Y, Yahiaoui N, Ruiz M, Santoni S, Labouisse JP, Pham JL, David J, Glémin S. 2017. Evolutionary forces affecting synonymous variations in plant genomes. PLoS Genetics. 13:1–28.

Copley SD. 2020, April. Evolution of new enzymes by gene duplication and divergence. The FEBS journal. 287(7):1262–1283.

Daron J, Bravo IG. 2021. Variability in codon usage in coronaviruses is mainly driven by mutational bias and selective constraints on cpg dinucleotide. Viruses. 13:1800.

Donizetti A, Fiengo M, Minucci S, Aniello F. 2009, October. Duplicated zebrafish relaxin-3 gene shows a different expression pattern from that of the co-orthologue gene. Development, Growth & Differentiation. 51(8):715–722.

Duret L. 2002, December. Evolution of synonymous codon usage in metazoans. Current Opinion in Genetics & Development. 12(6):640–649.

Duret L, Mouchiroud D. 1999, April. Expression pattern and, surprisingly, gene length shape codon usage in Caenorhabditis, Drosophila, and Arabidopsis. Proceedings of the National Academy of Sciences. 96(8):4482–4487. Publisher: National Academy of Sciences Section: Biological Sciences.

Ferris SD, Whitt GS. 1979, April. Evolution of the differential regulation of duplicate genes after polyploidization. Journal of Molecular Evolution. 12(4):267–317.

Freilich S, Massingham T, Blanc E, Goldovsky L, Thornton JM. 2006. Relating tissue specialization to the differenti-ation of expression of singleton and duplicate mouse proteins. Genome Biology. 7(10):R89.

Fu J, Dang Y, Counter C, Liu Y. 2018. Codon usage regulates human KRAS expression at both transcriptional and translational levels. The Journal of Biological Chemistry. 293(46):17929–17940.

Galtier N, Roux C, Rousselle M, Romiguier J, Figuet E, Glémin S, Bierne N, Duret L. 2018, May. Codon Usage Bias in Animals: Disentangling the Effects of Natural Selection, Effective Population Size, and GC-Biased Gene Conversion. Molecular Biology and Evolution. 35(5):1092–1103.

Grantham R, Gautier C, Gouy M, Mercier R, Pavé A. 1980, January. Codon catalog usage and the genome hypothesis. Nucleic Acids Research. 8(1):r49–r62.

Guschanski K, Warnefors M, Kaessmann H. 2017. The evolution of duplicate gene expression in mammalian organs. Genome Research. 27(9):1461–1474.

Hannigan MM, Zagore LL, Licatalosi DD. 2017, June. Ptbp2 controls an alternative splicing network required for cell communication during spermatogenesis. Cell reports. 19(12):2598–2612.

Hershberg R, Petrov DA. 2009, July. General rules for optimal codon choice. PLoS genetics. 5(7):e1000556.

Holmquist GP. 1989, June. Evolution of chromosome bands: Molecular ecology of noncoding DNA. Journal of Molecular Evolution. 28(6):469–486.

Ikemura T. 1981, September. Correlation between the abundance of Escherichia coli transfer RNAs and the occurrence of the respective codons in its protein genes: a proposal for a synonymous codon choice that is optimal for the E. coli translational system. Journal of Molecular Biology. 151(3):389–409.

Ivanov IP, Firth AE, Michel AM, Atkins JF, Baranov PV. 2011, May. Identification of evolutionarily conserved non-AUG-initiated N-terminal extensions in human coding sequences. Nucleic Acids Research. 39(10):4220–4234.

Katoh K, Misawa K, Kuma Ki, Miyata T. 2002, July. MAFFT: a novel method for rapid multiple sequence alignment based on fast Fourier transform. Nucleic Acids Research. 30(14):3059–3066.

Keppetipola N, Sharma S, Li Q, Black DL. 2012, August. Neuronal regulation of pre-mRNA splicing by polypyrim-idine tract binding proteins, PTBP1 and PTBP2. Critical Reviews in Biochemistry and Molecular Biology. 47(4):360–378.

Khorana HG, Büchi H, Ghosh H, Gupta N, Jacob TM, Kössel H, Morgan R, Narang SA, Ohtsuka E, Wells RD. 1966. Polynucleotide synthesis and the genetic code. Cold Spring Harbor Symposia on Quantitative Biology. 31:39–49.

Koonin EV. 2005. Orthologs, Paralogs, and Evolutionary Genomics. Annual Review of Genetics. 39(1):309–338. _eprint: https://doi.org/10.1146/annurev.genet.39.073003.114725.

Kumar S, Stecher G, Suleski M, Hedges SB. 2017. TimeTree: A Resource for Timelines, Timetrees, and Divergence Times. Molecular Biology and Evolution. 34(7):1812–1819.

Lampson BL, Pershing NLK, Prinz JA, Lacsina JR, Marzluff WF, Nicchitta CV, MacAlpine DM, Counter CM. 2013, January. Rare codons regulate KRas oncogenesis. Current biology: CB. 23(1):70–75.

Laurin-Lemay S, Rodrigue N, Lartillot N, Philippe H. 2018. Conditional Approximate Bayesian Computation: A New Approach for Across-Site Dependency in High-Dimensional Mutation-Selection Models. Molecular Biology and Evolution. 35(11):2819–2834.

Lujan SA, Williams JS, Pursell ZF, Abdulovic-Cui AA, Clark AB, McElhinny SAN, Kunkel TA. 2012, October. Mis-match Repair Balances Leading and Lagging Strand DNA Replication Fidelity. PLOS Genetics. 8(10):e1003016. Publisher: Public Library of Science.

Makeyev EV, Zhang J, Carrasco MA, Maniatis T. 2007, August. The MicroRNA miR-124 Promotes Neuronal Differentiation by Triggering Brain-Specific Alternative Pre-mRNA Splicing. Molecular cell. 27(3):435–448.

Marais G. 2003, June. Biased gene conversion: implications for genome and sex evolution. Trends in Genetics. 19(6):330–338. Publisher: Elsevier.

Merkin J, Russell C, Chen P, Burge CB. 2012, December. Evolutionary dynamics of gene and isoform regulation in Mammalian tissues. Science (New York, N.Y.). 338(6114):1593–1599.

Mordstein C, Savisaar R, Young RS, Bazile J, Talmane L, Luft J, Liss M, Taylor MS, Hurst LD, Kudla G. 2020, April. Codon Usage and Splicing Jointly Influence mRNA Localization. Cell Systems. 10(4):351–362.e8.

Munk M, Villalobo E, Villalobo A, Berchtold MW. 2022, November. Differential expression of the three independent cam genes coding for an identical protein: Potential relevance of distinct mrna stability by different codon usage. Cell Calcium. 107.

NCBI Resource Coordinators. 2018. Database resources of the National Center for Biotechnology Information. Nucleic Acids Research. 46(D1):D8–D13.

Newman ZR, Young JM, Ingolia NT, Barton GM. 2016, March. Differences in codon bias and GC content contribute to the balanced expression of TLR7 and TLR9. Proceedings of the National Academy of Sciences of the United States of America. 113(10):E1362–1371.

Nirenberg MW, Matthaei JH. 1961, October. The dependence of cell-free protein synthesis in e. coli upon naturally occurring or synthetic polyribonucleotides. Proceedings of the National Academy of Sciences of the United States of America. 47(10):1588–1602.

Novoa EM, Jungreis I, Jaillon O, Kellis M. 2019. Elucidation of Codon Usage Signatures across the Domains of Life. Molecular Biology and Evolution. 36(10):2328–2339.

Novoa EM, Ribas de Pouplana L. 2012, November. Speeding with control: codon usage, tRNAs, and ribosomes. Trends in genetics: TIG. 28(11):574–581.

Palidwor GA, Perkins TJ, Xia X. 2010, October. A general model of codon bias due to GC mutational bias. PloS One. 5(10):e13431.

Peabody DS. 1989, March. Translation initiation at non-AUG triplets in mammalian cells. The Journal of Biological Chemistry. 264(9):5031–5035.

Percudani R, Pavesi A, Ottonello S. 1997, May. Transfer RNA gene redundancy and translational selection in Saccha-romyces cerevisiae11Edited by J. Karn. Journal of Molecular Biology. 268(2):322–330.

Pina J, Ontiveros RJ, Keppetipola N, Nikolaidis N. 2018, April. A Bioinformatics Approach to Discover the Evolutionary Origin of the PTBP Splicing Regulators. The FASEB Journal. 32(1_supplement):802.16–802.16. Publisher: Federation of American Societies for Experimental Biology.

Plotkin JB, Kudla G. 2011, January. Synonymous but not the same: the causes and consequences of codon bias. Nature Reviews Genetics. 12(1):32–42.

Pouyet F, Mouchiroud D, Duret L, Sémon M. 2017. Recombination, meiotic expression and human codon usage. eLife. 6.

Presnyak V, Alhusaini N, Chen YH, Martin S, Morris N, Kline N, Olson S, Weinberg D, Baker KE, Graveley BR, Coller J. 2015, March. Codon optimality is a major determinant of mRNA stability. Cell. 160(6):1111–1124.

Reijns MAM, Kemp H, Ding J, Marion de Procé S, Jackson AP, Taylor MS. 2015, February. Lagging-strand replication shapes the mutational landscape of the genome. Nature. 518(7540):502–506. Number: 7540 Publisher: Nature Publishing Group.

Robinson DF, Foulds LR. 1981, February. Comparison of phylogenetic trees. Mathematical Biosciences. 53(1):131–147.

Robinson F, Jackson RJ, Smith CWJ. 2008, March. Expression of Human nPTB Is Limited by Extreme Suboptimal Codon Content. PLOS ONE. 3(3):e1801. Publisher: Public Library of Science.

Satapathy SS, Powdel BR, Buragohain AK, Ray SK. 2016, October. Discrepancy among the synonymous codons with respect to their selection as optimal codon in bacteria. DNA Research. 23(5):441–449. Publisher: Oxford Academic.

Scornavacca C, Belkhir K, Lopez J, Dernat R, Delsuc F, Douzery EJP, Ranwez V. 2019, April. OrthoMaM v10: Scaling-Up Orthologous Coding Sequence and Exon Alignments with More than One Hundred Mammalian Genomes. Molecular Biology and Evolution. 36(4):861–862. Publisher: Oxford Academic.

Sharp PM, Li WH. 1987. The codon Adaptation Index–a measure of directional synonymous codon usage bias, and its potential applications. Nucleic Acids Research. 15(3):1281–1295.

Sonnhammer ELL, Koonin EV. 2002, December. Orthology, paralogy and proposed classification for paralog subtypes. Trends in genetics: TIG. 18(12):619–620.

Soria-Carrasco V, Talavera G, Igea J, Castresana J. 2007, November. The K tree score: quantification of differences in the relative branch length and topology of phylogenetic trees. Bioinformatics (Oxford, England). 23(21):2954–2956.

Spellman R, Llorian M, Smith CW. 2007, August. Crossregulation and functional redundancy between the splicing regulator ptb and its paralogs nptb and rod1. Molecular Cell. 27:420–434.

Spencer PS, Barral JM. 2012, March. Genetic code redundancy and its influence on the encoded polypeptides. Computational and Structural Biotechnology Journal. 1.

Stamatakis A. 2014, May. RAxML version 8: a tool for phylogenetic analysis and post-analysis of large phylogenies. Bioinformatics (Oxford, England). 30(9):1312–1313.

Vuong JK, Lin CH, Zhang M, Chen L, Black DL, Zheng S. 2016. PTBP1 and PTBP2 Serve Both Specific and Redundant Functions in Neuronal Pre-mRNA Splicing. Cell Reports. 17(10):2766–2775.

Wagner EJ, Garcia-Blanco MA. 2002, October. Rnai-mediated ptb depletion leads to enhanced exon definition. Molecular Cell. 10:943–949.

Whittle CA, Extavour CG. 2016, September. Expression-Linked Patterns of Codon Usage, Amino Acid Frequency, and Protein Length in the Basally Branching Arthropod Parasteatoda tepidariorum. Genome Biology and Evolution. 8(9):2722–2736. Publisher: Oxford Academic.

Yates AD, Achuthan P, Akanni W, Allen J, Allen J, Alvarez-Jarreta J, Amode MR, Armean IM, Azov AG, Bennett R, Bhai J, Billis K, Boddu S, Marugán JC, Cummins C, Davidson C, Dodiya K, Fatima R, Gall A, Giron CG, Gil L, Grego T, Haggerty L, Haskell E, Hourlier T, Izuogu OG, Janacek SH, Juettemann T, Kay M, Lavidas I, Le T, Lemos D, Martinez JG, Maurel T, McDowall M, McMahon A, Mohanan S, Moore B, Nuhn M, Oheh DN, Parker A, Parton A, Patricio M, Sakthivel MP, Abdul Salam AI, Schmitt BM, Schuilenburg H, Sheppard D, Sycheva M, Szuba M, Taylor K, Thormann A, Threadgold G, Vullo A, Walts B, Winterbottom A, Zadissa A, Chakiachvili M, Flint B, Frankish A, Hunt SE, IIsley G, Kostadima M, Langridge N, Loveland JE, Martin FJ, Morales J, Mudge JM, Muffato M, Perry E, Ruffier M, Trevanion SJ, Cunningham F, Howe KL, Zerbino DR, Flicek P. 2020, January. Ensembl 2020. Nucleic Acids Research. 48(D1):D682–D688. Publisher: Oxford Academic.

Yu Y, Fuscoe JC, Zhao C, Guo C, Jia M, Qing T, Bannon DI, Lancashire L, Bao W, Du T, Luo H, Su Z, Jones WD, Moland CL, Branham WS, Qian F, Ning B, Li Y, Hong H, Guo L, Mei N, Shi T, Wang KY, Wolfinger RD, Nikolsky Y, Walker SJ, Duerksen-Hughes P, Mason CE, Tong W, Thierry-Mieg J, Thierry-Mieg D, Shi L, Wang C. 2014, February. A rat RNA-Seq transcriptomic BodyMap across 11 organs and 4 developmental stages. Nature Communications. 5(1):3230.Number: 1 Publisher: Nature Publishing Group.

Zagore LL, Grabinski SE, Sweet TJ, Hannigan MM, Sramkoski RM, Li Q, Licatalosi DD. 2015, December. RNA Binding Protein Ptbp2 Is Essential for Male Germ Cell Development. Molecular and Cellular Biology. 35(23):4030–4042.

