## Supplementary material for "Subfunctionalisation of paralogous genes and evolution of differential codon usage preferences: the showcase of polypyrimidine tract binding proteins"

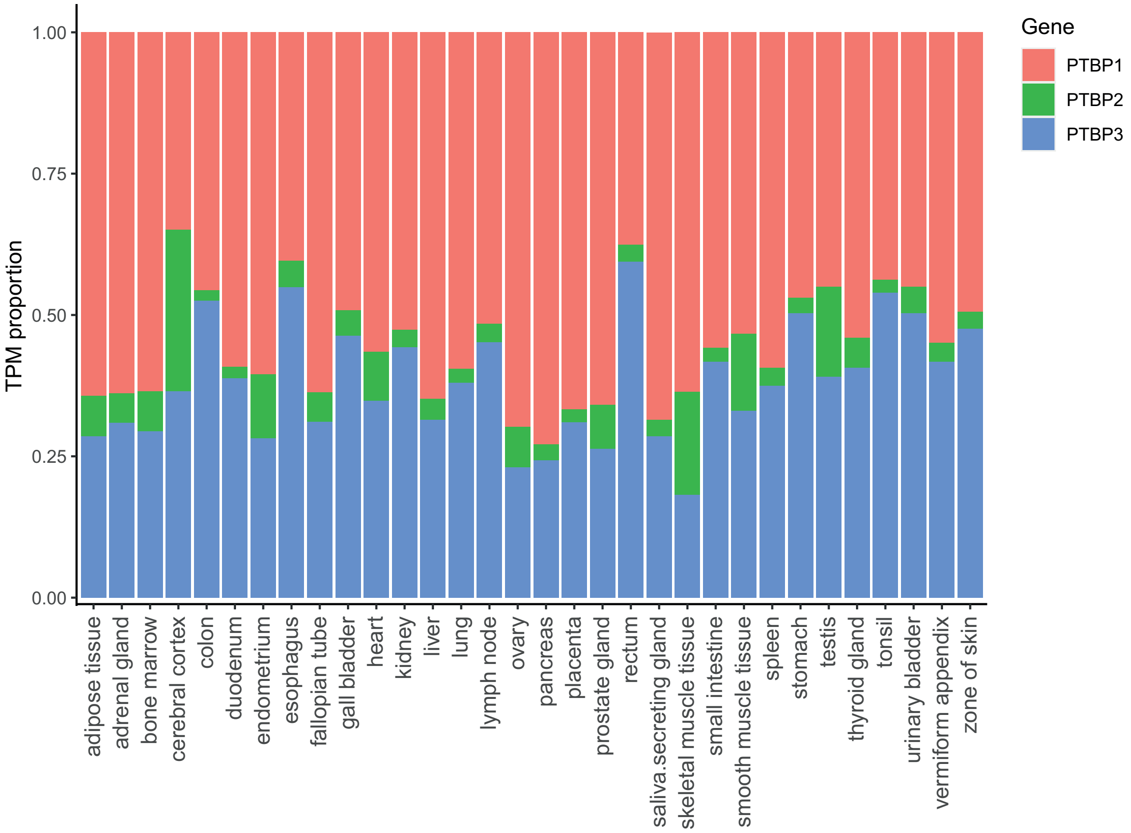


Figure S1: **Relative mRNA levels of *PTBP1* (red), *PTBP2* (green) and *PTBP3* (blue) in 32 human tissues.** Values given in this graph represent the proportion of transcripts per million (TPM) for each *PTBP*, as communicated by Zhang and coworkers (Zhang et al., 2012) . Overall *PTBP1* is the most transcribed gene followed by *PTBP3*, while *PTBP2* shows maximum expression in cerebral cortex and in skeletal muscle.

| | *PTBP1* | | | | *PTBP2* | | | | *PTBP3* | | | |
|---|---|---|---|---|---|---|---|---|---|---|---|---|
| **Species** | **%GC** | **%GC1** | **%GC2** | **%GC3** | **%GC** | **%GC1** | **%GC2** | **%GC3** | **%GC** | **%GC1** | **%GC2** | **%GC3** |
| **Mammalians** | **59.71** | **58.16** | **41.19** | **79.77** | **41.88** | **55.72** | **38.25** | **31.66** | **44.51** | **56.30** | **41.01** | **36.23** |
| *Aotus nancymaae* | 61.33 | 58.29 | 40.76 | 84.95 | 41.20 | 55.56 | 38.29 | 29.76 | 44.25 | 55.94 | 41.57 | 35.25 |
| *Balaenoptera acutorostrata* | 61.65 | 58.10 | 41.14 | 85.71 | 41.20 | 55.95 | 38.29 | 29.37 | 43.81 | 56.90 | 41.00 | 33.53 |
| *Bos taurus* | 59.30 | 58.29 | 41.52 | 78.10 | 41.53 | 55.75 | 38.29 | 30.56 | 44.87 | 56.79 | 40.92 | 36.90 |
| *Canis lupus* | 60.00 | 58.48 | 40.95 | 80.57 | 41.53 | 55.75 | 38.29 | 30.56 | 44.49 | 56.57 | 41.10 | 35.81 |
| *Cebus capucinus* | 61.19 | 57.92 | 39.08 | 86.57 | 41.47 | 55.56 | 38.29 | 30.56 | 44.19 | 56.13 | 41.38 | 35.06 |
| *Ceratotherium simum* | 62.35 | 58.67 | 41.33 | 87.05 | 41.07 | 55.36 | 38.29 | 29.56 | 44.57 | 56.46 | 40.66 | 36.61 |
| *Chinchilla lanigera* | 58.92 | 57.52 | 41.33 | 77.91 | 42.06 | 55.75 | 38.29 | 32.14 | 44.72 | 56.43 | 40.88 | 36.85 |
| *Chrysochloris asiatica* | 58.86 | 58.29 | 41.33 | 76.95 | 40.62 | 55.31 | 38.08 | 28.46 | 43.97 | 55.39 | 40.96 | 35.58 |
| *Condylura cristata* | 61.08 | 57.71 | 41.33 | 84.19 | 41.34 | 55.95 | 38.29 | 29.76 | 43.31 | 55.47 | 40.69 | 33.78 |
| *Dasypus novemcinctus* | 65.08 | 61.17 | 43.41 | 90.66 | 40.87 | 55.36 | 38.29 | 28.97 | 44.06 | 55.75 | 41.00 | 35.44 |
| *Delphinapterus leucas* | 61.33 | 57.71 | 41.33 | 84.95 | 41.15 | 55.51 | 38.08 | 29.86 | 44.19 | 57.28 | 41.19 | 34.10 |
| *Desmodus rotundus* | 56.44 | 57.91 | 40.95 | 70.48 | 51.98 | 57.94 | 38.49 | 59.52 | 43.61 | 55.75 | 40.81 | 34.29 |
| *Echinops telfairi* | 60.96 | 59.89 | 41.07 | 81.94 | 40.99 | 55.29 | 37.92 | 29.74 | 44.96 | 56.51 | 41.00 | 37.36 |
| *Equus asinus* | 63.55 | 58.59 | 41.22 | 90.84 | 41.40 | 55.56 | 38.29 | 30.36 | 45.86 | 56.84 | 40.46 | 40.27 |
| *Equus caballus* | 63.61 | 58.78 | 41.22 | 90.84 | 41.67 | 55.56 | 38.29 | 31.15 | 46.05 | 56.84 | 40.46 | 40.85 |
| *Equus przewalskii* | 63.15 | 57.39 | 41.46 | 90.60 | 41.67 | 55.56 | 38.29 | 31.15 | 46.05 | 56.84 | 40.46 | 40.85 |
| *Erinaceus europaeus* | 63.80 | 58.78 | 40.84 | 91.79 | 40.34 | 55.75 | 37.90 | 27.38 | 43.18 | 54.58 | 41.34 | 33.61 |
| *Felis catus* | 60.64 | 58.48 | 40.95 | 82.48 | 41.27 | 55.75 | 38.29 | 29.76 | 44.36 | 56.15 | 40.96 | 35.96 |
| *Heterocephalus glaber* | 59.75 | 57.91 | 41.52 | 79.81 | 41.47 | 55.75 | 38.29 | 30.36 | 43.81 | 56.32 | 40.81 | 34.29 |
| *Homo sapiens* | 61.27 | 58.67 | 40.76 | 84.38 | 41.07 | 55.56 | 38.29 | 29.37 | 43.93 | 56.07 | 41.23 | 34.49 |
| *Ictidomys tridecemlineatus* | 60.06 | 57.52 | 41.33 | 81.33 | 41.20 | 55.75 | 38.29 | 29.56 | 43.17 | 55.36 | 41.00 | 33.14 |
| *Loxodonta africana* | 58.90 | 58.19 | 40.66 | 77.84 | 41.07 | 55.75 | 38.29 | 29.17 | 44.15 | 56.24 | 41.08 | 35.13 |
| *Macaca fascicularis* | 60.69 | 58.40 | 40.65 | 83.02 | 41.27 | 55.56 | 38.29 | 29.96 | 43.93 | 56.32 | 41.19 | 34.29 |
| *Microcebus murinus* | 59.81 | 57.52 | 41.14 | 80.76 | 40.41 | 55.16 | 38.29 | 27.78 | 43.30 | 55.56 | 41.00 | 33.33 |
| *Miniopterus natalensis* | 55.05 | 57.52 | 40.95 | 66.67 | 47.82 | 56.75 | 38.29 | 48.41 | 45.50 | 58.02 | 40.88 | 37.58 |
| *Monodelphis domestica* | 46.98 | 55.43 | 41.14 | 44.38 | 41.01 | 54.96 | 38.29 | 29.76 | 43.35 | 55.68 | 41.23 | 33.14 |
| *Mus musculus* | 57.17 | 57.94 | 41.49 | 72.08 | 43.59 | 56.35 | 38.10 | 36.31 | 46.64 | 56.05 | 41.08 | 42.80 |
| *Ochotona princeps* | 61.27 | 58.86 | 41.52 | 83.43 | 41.93 | 55.75 | 38.29 | 31.75 | 46.11 | 57.09 | 41.19 | 40.04 |
| *Odocoileus virginianus* | 59.24 | 58.48 | 41.52 | 77.71 | 41.73 | 55.75 | 38.29 | 31.15 | 46.25 | 57.44 | 41.60 | 39.70 |
| *Orcinus orca* | 61.14 | 57.91 | 41.33 | 84.19 | 41.27 | 55.95 | 38.29 | 29.56 | 44.19 | 57.09 | 41.19 | 34.29 |
| *Ornithorhynchus anatinus* | 49.27 | 55.62 | 41.14 | 51.05 | 48.63 | 56.11 | 37.88 | 51.90 | 44.89 | 52.60 | 39.88 | 42.20 |
| *Orycteropus afer* | 60.00 | 59.62 | 41.33 | 79.05 | 40.75 | 54.91 | 38.08 | 29.26 | 45.34 | 55.94 | 41.38 | 38.70 |
| *Oryctolagus cuniculus* | - | - | - | - | 41.53 | 55.16 | 38.10 | 31.35 | 45.79 | 56.32 | 41.00 | 40.04 |
| *Ovis aries* | 59.24 | 58.48 | 41.52 | 77.71 | 41.55 | 55.87 | 38.37 | 30.42 | 45.04 | 57.06 | 41.41 | 36.64 |
| *Pan troglodytes* | 61.27 | 58.67 | 40.76 | 84.38 | 41.20 | 55.56 | 38.29 | 29.76 | 43.93 | 56.13 | 41.19 | 34.48 |
| *Panthera pardus* | 61.08 | 58.29 | 40.95 | 84.00 | 41.20 | 55.75 | 38.29 | 29.56 | 44.34 | 56.05 | 40.88 | 36.08 |
| *Pantholops hodgsonii* | 59.30 | 58.48 | 41.52 | 77.91 | 41.47 | 56.35 | 38.49 | 29.56 | 45.72 | 58.19 | 41.37 | 37.61 |
| *Papio anubis* | 60.62 | 58.40 | 40.65 | 82.82 | 41.07 | 55.36 | 38.29 | 29.56 | 44.06 | 56.32 | 41.19 | 34.67 |
| *Phascolarctos cinereus* | 47.87 | 55.05 | 41.14 | 47.43 | 40.87 | 55.36 | 38.29 | 28.97 | 43.96 | 55.56 | 42.14 | 34.17 |
| *Physeter catodon* | 61.27 | 57.71 | 40.95 | 85.14 | 41.49 | 56.28 | 38.29 | 29.92 | 43.87 | 57.09 | 41.00 | 33.53 |
| *Pteropus vampyrus* | 63.11 | 58.86 | 41.33 | 89.14 | 42.20 | 55.95 | 38.29 | 32.34 | 43.95 | 56.24 | 41.08 | 34.55 |
| *Rousettus aegyptiacus* | 63.18 | 58.86 | 41.33 | 89.33 | 42.13 | 55.95 | 38.29 | 32.14 | 44.15 | 56.24 | 41.08 | 35.13 |
| *Sarcophilus harrisii* | 46.98 | 54.48 | 41.14 | 45.33 | 41.01 | 55.36 | 38.29 | 29.37 | 43.48 | 55.88 | 40.66 | 33.91 |
| *Sorex araneus* | 64.34 | 59.89 | 41.62 | 91.53 | 42.73 | 56.55 | 38.29 | 33.33 | 45.87 | 56.81 | 40.69 | 40.12 |
| *Trichechus manatus* | 61.33 | 59.05 | 41.52 | 83.43 | 41.02 | 55.11 | 38.08 | 29.86 | 44.63 | 56.10 | 40.98 | 36.79 |
| *Tupaia belangeri* | 63.30 | 58.97 | 41.60 | 89.31 | 40.74 | 55.16 | 38.29 | 28.77 | 44.40 | 57.80 | 40.22 | 35.17 |
| *Zalophus californianus* | 59.87 | 58.67 | 41.14 | 79.81 | 41.34 | 55.75 | 38.29 | 29.96 | 43.95 | 56.05 | 41.08 | 34.74 |
| **Non-mammalians** | **49.42** | **55.61** | **40.81** | **51.84** | **43.19** | **54.93** | **38.48** | **36.15** | **45.77** | **55.05** | **40.73** | **41.53** |
| *Acanthisitta chloris* | 54.52 | 56.30 | 41.99 | 65.27 | 42.94 | 55.47 | 38.37 | 34.99 | - | - | - | - |
| *Alligator mississippiensis* | 47.37 | 55.43 | 40.95 | 45.71 | 42.21 | 55.07 | 38.17 | 33.40 | 45.34 | 55.88 | 41.23 | 38.92 |
| *Anas platyrhynchos* | 47.11 | 54.67 | 40.76 | 45.91 | 43.25 | 55.75 | 38.29 | 35.71 | 44.86 | 56.45 | 40.43 | 37.70 |
| *Anser cygnoides* | 47.11 | 54.67 | 40.76 | 45.91 | 44.30 | 56.20 | 41.92 | 34.77 | 44.70 | 54.91 | 40.66 | 38.54 |
| *Apteryx australis* | 47.37 | 55.62 | 40.76 | 45.71 | 42.53 | 55.95 | 38.29 | 33.33 | 44.70 | 54.79 | 40.61 | 38.70 |
| *Chaetura pelagica* | 50.00 | 56.01 | 40.31 | 53.68 | 42.69 | 55.51 | 38.28 | 34.27 | 45.20 | 54.67 | 41.44 | 39.49 |
| *Chrysemys picta* | 48.22 | 54.77 | 41.03 | 48.86 | 42.20 | 54.96 | 38.29 | 33.33 | 45.15 | 55.49 | 40.85 | 39.11 |
| *Crocodylus porosus* | 46.79 | 55.24 | 40.95 | 44.19 | 42.13 | 54.76 | 38.29 | 33.33 | 45.28 | 56.26 | 41.23 | 38.34 |
| *Danio rerio* | 50.13 | 55.15 | 41.60 | 53.63 | 51.35 | 55.05 | 40.20 | 58.81 | 52.24 | 55.77 | 40.96 | 60.00 |

| | | | | | | | | | | | | |
|---|---|---|---|---|---|---|---|---|---|---|---|---|
| *Gallus gallus* | 49.18 | 56.25 | 40.53 | 50.76 | 42.13 | 55.56 | 38.29 | 32.54 | 43.18 | 52.30 | 39.12 | 38.12 |
| *Gavialis gangeticus* | 47.05 | 55.43 | 40.95 | 44.76 | 42.02 | 54.87 | 38.17 | 33.00 | 45.34 | 56.26 | 41.23 | 38.54 |
| *Gekko japonicus* | 53.79 | 55.26 | 40.15 | 65.97 | 41.62 | 55.07 | 38.17 | 31.61 | 45.34 | 54.72 | 41.04 | 40.27 |
| *Haliaeetus albicilla* | 46.73 | 55.05 | 40.95 | 44.19 | 42.73 | 55.36 | 38.10 | 34.72 | 44.36 | 53.39 | 40.62 | 39.07 |
| *Latimeria chalumnae* | 45.87 | 54.13 | 40.88 | 42.61 | 42.59 | 55.36 | 38.49 | 33.93 | 46.69 | 56.26 | 41.04 | 42.78 |
| *Lepisosteus oculatus* | 50.45 | 55.60 | 40.35 | 55.41 | 45.67 | 54.65 | 39.73 | 42.64 | 51.47 | 55.00 | 40.77 | 58.65 |
| *Nothoprocta perdicaria* | 48.77 | 56.17 | 40.61 | 49.53 | 44.09 | 55.51 | 38.08 | 38.68 | 44.64 | 53.95 | 41.04 | 38.92 |
| *Numida meleagris* | 49.62 | 55.92 | 40.65 | 52.29 | 42.79 | 56.35 | 38.29 | 33.73 | 43.31 | 53.09 | 39.72 | 37.13 |
| *Oncorhynchus mykiss* | 57.55 | 56.79 | 39.62 | 76.23 | 54.36 | 54.56 | 39.55 | 68.97 | 55.74 | 55.92 | 43.80 | 67.49 |
| *Phasianus colchicus* | 49.05 | 56.30 | 40.46 | 50.38 | 42.15 | 55.87 | 38.17 | 32.41 | 42.29 | 52.31 | 38.08 | 36.47 |
| *Podarcis muralis* | 55.11 | 55.24 | 41.14 | 68.95 | 42.39 | 54.37 | 38.10 | 34.72 | 45.05 | 55.02 | 40.54 | 39.58 |
| *Pogona vitticeps* | 60.71 | 57.22 | 41.26 | 83.65 | 41.53 | 54.96 | 37.90 | 31.75 | 44.57 | 55.49 | 41.43 | 36.80 |
| *Python bivittatus* | 47.79 | 55.09 | 40.88 | 47.41 | 40.01 | 53.11 | 37.88 | 29.06 | 44.77 | 55.11 | 41.81 | 37.38 |
| *Rhinatrema bivittatum* | 48.47 | 55.73 | 41.41 | 48.28 | 41.27 | 54.17 | 38.49 | 31.15 | 44.99 | 55.27 | 40.82 | 38.87 |
| *Rhincodon typus* | 46.63 | 54.20 | 41.41 | 44.28 | 43.19 | 52.34 | 38.52 | 38.72 | 45.65 | 56.78 | 37.72 | 42.44 |
| *Struthio camelus* | 46.92 | 55.24 | 40.76 | 44.76 | 42.86 | 55.75 | 38.29 | 34.52 | 45.73 | 56.26 | 41.81 | 39.11 |
| *Xenopus laevis* | 45.89 | 57.28 | 40.19 | 40.19 | 40.76 | 53.41 | 37.35 | 31.53 | 44.68 | 55.13 | 40.23 | 38.69 |
| *Xenopus tropicalis* | 46.10 | 56.67 | 40.43 | 41.20 | 40.30 | 53.21 | 37.35 | 30.32 | 44.68 | 54.74 | 40.62 | 38.69 |

| **Protostoma *PTBP*** | | | | |
|---|---|---|---|---|
| | **%GC** | **%GC1** | **%GC2** | **%GC3** |
| *Apis mellifera* | 51.36 | 54.07 | 38.70 | 61.30 |
| *Crassostrea gigas* | 52.66 | 57.14 | 41.03 | 59.80 |
| *Drosophila melanogaster* | 53.84 | 54.11 | 41.12 | 66.28 |

Table S2: **Percentages of total GC content and GC content at the first (GC1), second (GC2) and third (GC3) position of nucleotides for *PTBP1*, *PTBP2* and *PTBP3***. Lines in bold present the mean score values for mammalians and non-mammalians species.

Table S3: **Exomic (third position), intronic and flanking regions GC content of *PTBP1*, *PTBP2* and *PTBP3* for the fifteen species used in the genomic context analysis**. For each species, the assembly number to extract the corresponding values is given in parenthesis.

| | *PTBP1* GC content (%) | | | *PTBP2* GC content (%) | | | *PTBP3* GC content (%) | | |
|---|---|---|---|---|---|---|---|---|---|
| Species (Assembly number) | GC3 | Introns | Flanks | GC3 | Introns | Flanks | GC3 | Introns | Flanks |
| *Bos taurus* (6369068) | 77.1 | 60.1 | 52.1 | 31. | 34.6 | 38.3 | 37.7 | 36.1 | 40.7 |
| *Canis familiaris* (313658) | 81.8 | 62.5 | 68.4 | 31.2 | 32.7 | 35.8 | 36.1 | 35.8 | 41.5 |
| *Danio rerio* (482478 ) | 53.0 | 35.2 | 34.8 | 57.7 | 36.7 | 36.7 | 60.2 | 33.9 | 33.6 |
| *Dasypus novemcinctus* (326198) | 88.6 | 67.6 | 51.3 | 29.5 | 33.6 | 36.3 | 35.3 | 36.3 | 37.2 |
| *Equus caballus* (286568) | 85.0 | 66.9 | 66.5 | 32.1 | 34.2 | 37.1 | 46.6 | 37.12 | 42.2 |
| *Gallus gallus* (6347868) | 50.1 | 42.6 | 53.1 | 33.3 | 36.4 | 39.9 | 37.3 | 36.4 | 48.3 |
| *Gekko japonicus* (2693898) | 40.4 | 42.3 | 43.8 | - | - | - | 64.6 | 47.0 | 49.7 |
| *Homo sapiens* (8687898) | 82.4 | 60.1 | 57.3 | 30.8 | 35.7 | 37.1 | 36.1 | 37.2 | 42.2 |
| *Latimeria chalumnae* (303548) | 42.2 | 34.0 | 32.9 | 31.6 | 32.8 | 35.0 | 42.639 | 36.5 | 38.4 |
| *Loxodonta africana* (3288) | - | - | - | 29.6 | 35.2 | 38.0 | 35.3 | 36.3 | 38.6 |
| *Mus musculus* (1700338) | 72.5 | 55.7 | 50.9 | 37.4 | 35.7 | 40.7 | 42.2 | 38.0 | 43.6 |
| *Pan troglodytes* (5907448) | 82.4 | 60.1 | 57.0 | 31.1 | 35.0 | 36.9 | 36.1 | 38.3 | 42.7 |
| *Pteropus vampyrus* (1410408) | 89.9 | 65.6 | 52.0 | 32.5 | 32.2 | 34.9 | 34.4 | 32.4 | 36.9 |
| *Rhincodon typus* (4170658) | - | - | - | 37.7 | 39.7 | 41.0 | - | - | - |
| *Xenopus tropicalis* (3341798) | 40.6 | 35.8 | 39.1 | 31.6 | 35.2 | 39.1 | 39.0 | 37.5 | 37.7 |

Table S4: $COUSIN_{59}$ **and** $CAI$ **scores of genomic context analysis species.** The reference used is the global CUPrefs of the organism, estimated using all CDSs above 100 amino acids in length.Here, since we calculate the CUPrefs of *PTBP*s against a reference dataset without performing any genomic context analysis, we replaced missing data by individuals *PTBP*s

| | ***PTBP1*** | | ***PTBP2*** | | ***PTBP3*** | |
|---|---|---|---|---|---|---|
| **Species** | $COUSIN_{59}$ | $CAI$ | $COUSIN_{59}$ | $CAI$ | $COUSIN_{59}$ | $CAI$ |
| *Bos taurus* | 1.785 | 0.787 | -0.534 | 0.618 | -0.323 | 0.642 |
| *Canis familiaris* | 1.976 | 0.821 | -0.499 | 0.664 | -0.207 | 0.698 |
| *Danio rerio* | 0.655 | 0.782 | 1.136 | 0.774 | 1.148 | 0.789 |
| *Dasypus novemncinctus* | 1.958 | 0.818 | -0.57 | 0.659 | -0.277 | 0.699 |
| *Equus caballus* | 1.831 | 0.813 | -0.489 | 0.652 | 0.107 | 0.698 |
| *Gallus gallus* | 0.764 | 0.769 | -0.006 | 0.731 | 0.327 | 0.745 |
| *Gekko japonicus* | 1.582 | 0.835 | -0.627 | 0.751 | 0.199 | 0.785 |
| *Homo sapiens* | 1.747 | 0.815 | -0.477 | 0.678 | -0.235 | 0.709 |
| *Latimeria chalumnae* | 0.728 | 0.818 | 1.218 | 0.824 | 0.4 | 0.808 |
| *Loxodonta africana* | 2.041 | 0.821 | -0.442 | 0.704 | -0.159 | 0.74 |
| *Mus musculus* | 1.842 | 0.829 | 0.017 | 0.708 | 0.154 | 0.73 |
| *Pan troglodytes* | 1.82 | 0.81 | -0.476 | 0.682 | -0.219 | 0.712 |
| *Pteropus vampyrus* | 2.112 | 0.84 | -0.427 | 0.683 | -0.287 | 0.703 |
| *Rhincodon typus* | 0.882 | 0.791 | 1.598 | 0.82 | 1.163 | 0.804 |
| *Xenopus tropicalis* | 1.347 | 0.817 | 1.559 | 0.806 | 1.264 | 0.808 |

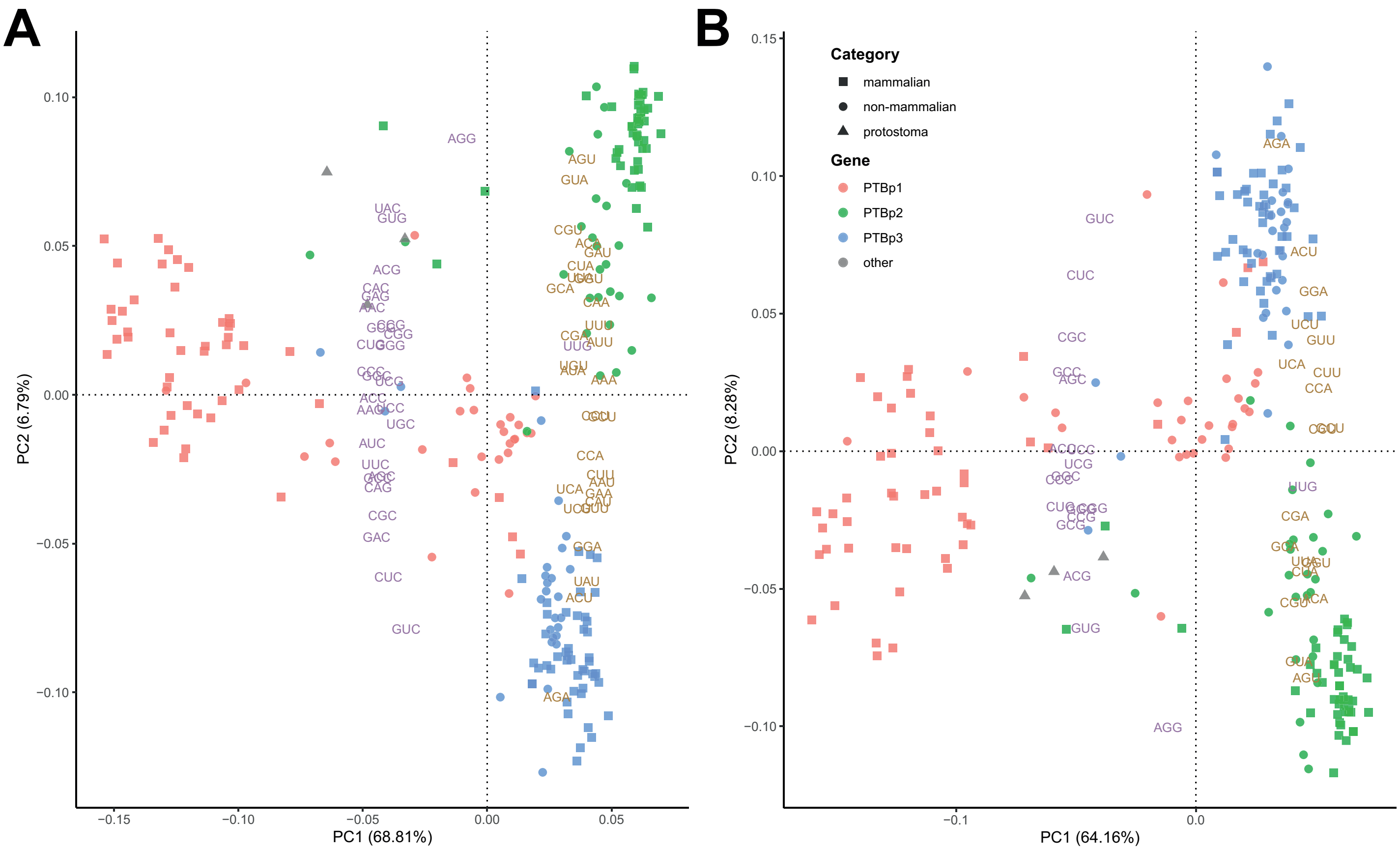


Figure S5: **Plot of the two first dimensions of a PCA analysis based on the codon usage preferences of A) all codons B) amino-acids encoded by four codons in *PTBP1*s (red), *PTBP2*s (green), *PTBP3*s (blue) and protostoma (grey) individuals.** Taxonomic information is included as mammals (squares), non-mammals (circles) and protostomates (triangles). The PCA was created using as variables the vectors of 59 positions (representing the relative frequencies of the 59 synonymous codons) for each individual gene. The eigenvalues of the individual codon variables are given by their position on the graph. Each codon variable is identified by its name and by a colour code, purple for GC-ending codons and orange for AT-ending codons. Note the position of the UUG-Leu codon, the sole GC-ending codon clustering with all other AT-ending codons in both panels. The percentage of the total variance explained by each axis is shown in parenthesis.

Table S6: **Comparison between species tree and subtrees of the nucleotide based maximum likelihood tree.** Each subtree corresponds to a paralog. The K-score compares topological and pairwise distances between trees, while the Robinson-Foulds score compares only topological distances between trees. As the K-score distance is not metric, the analysis has been run both ways.

| **Reference tree** | **Comparison tree** | **K-score** | **Scale factor** | **Robinson-Foulds** |
|---|---|---|---|---|
| **Nucleotide tree VS species tree** | | | | |
| PTBP1 | Species-1 | 0.759 | 0.001 | 42 |
| Species-1 | PTBP1 | 604.403 | 675.598 | 42 |
| PTBP2 | Species-2 | 0.762 | 0.001 | 24 |
| Species-2 | PTBP2 | 714.262 | 731.935 | 24 |
| PTBP3 | Species-3 | 1.700 | 0.001 | 28 |
| Species-3 | PTBP3 | 883.463 | 328.799 | 28 |
| **Nucleotide tree VS Amino acid tree** | | | | |
| PTBP1-AA | PTBP1-NT | 0.149 | 0.197 | 78 |
| PTBP1-NT | PTBP1-AA | 0.666 | 3.972 | 78 |
| PTBP2-AA | PTBP2-NT | 0.129 | 0.199 | 110 |
| PTBP2-NT | PTBP2-AA | 0.574 | 3.917 | 110 |
| PTBP3-AA | PTBP3-NT | 0.380 | 0.372 | 40 |
| PTBP3-NT | PTBP3-AA | 0.926 | 2.21 | 40 |

Table S7: **Results of a series of Mantel tests assessing correlation between pairwise nucleotide-based and amino acid-based distance matrices (AA-tree VS NT-tree) and between pairwise nucleotide-based and CUPrefs-based distance matrices (CUPrefs-tree VS NT-tree).** First part is comparison by gene of nucleotide and amino acid based maximum likelihood trees, second part is comparing the same nucleotide based trees by gene against pairwise distances of CUPrefs. Please note the different scales used for the different genes.

| **AA-tree VS NT-tree** | **Observation** | **simulated *p* value** |
|---|---|---|
| **Mammal** | | |
| PTBP1 | 0.712 | 0.001 |
| PTBP2 | 0.558 | 0.001 |
| PTBP3 | 0.987 | 0.001 |
| **Non-mammal** | | |
| PTBP1 | 0.929 | 0.001 |
| PTBP2 | 0.982 | 0.001 |
| PTBP3 | 0.748 | 0.001 |

| **CUPrefs VS NT-tree** | **Observation** | **simulated *p* value** |
|---|---|---|
| **Mammal** | | |
| PTBP1 | 0.908 | 0.001 |
| PTBP2 | 0.046 | 0.274 |
| PTBP3 | 0.737 | 0.001 |
| **Non-mammal** | | |
| PTBP1 | 0.091 | 0.273 |
| PTBP3 | -0.121 | 0.73 |
| PTBP3 | 0.145 | 0.165 |

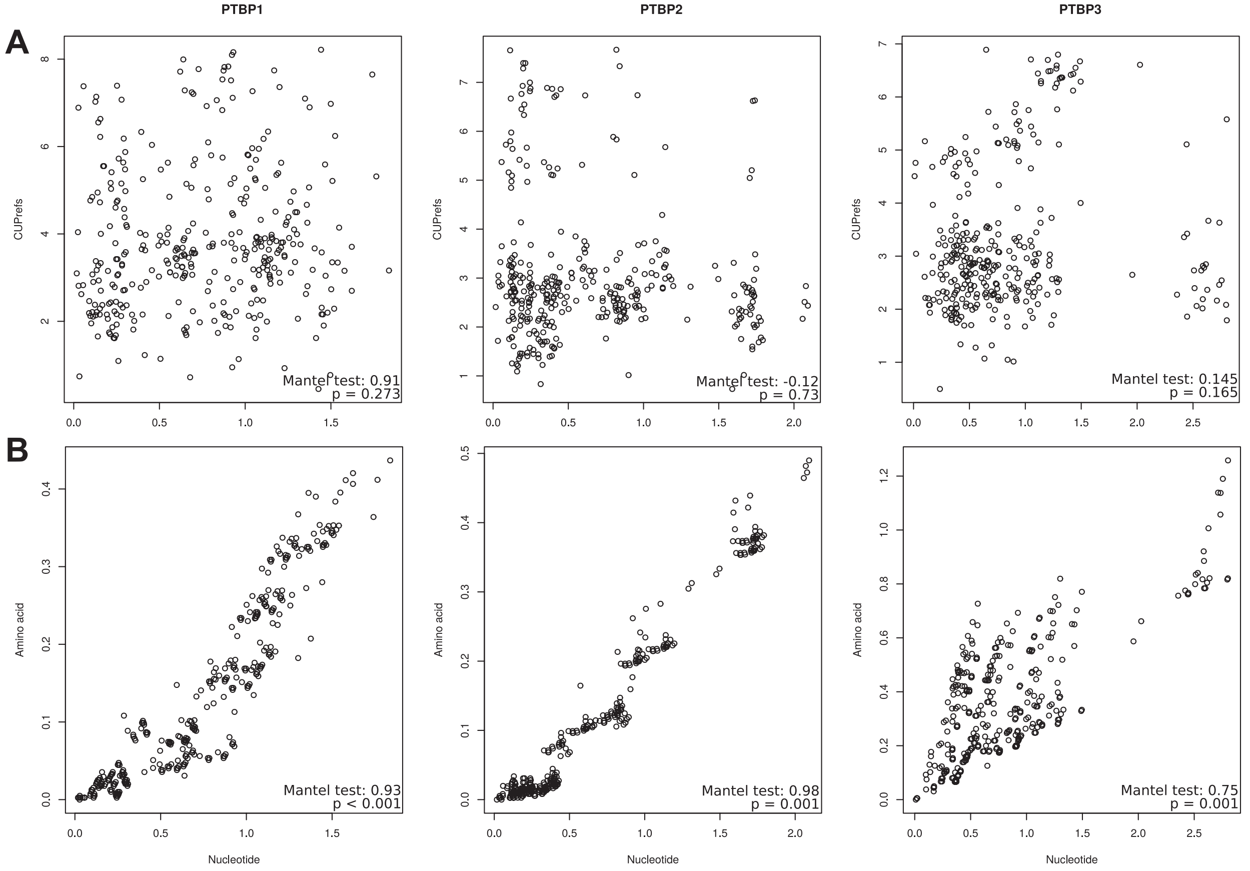


Figure S8: **Nucleotide-based pairwise distances against A) CUPrefs and B) amino-acid based pairwise distances for the different non-mammalian *PTBP* orthologs.** The results for a Mantel test assessing the correlation between the corresponding matrices are shown in the inset.

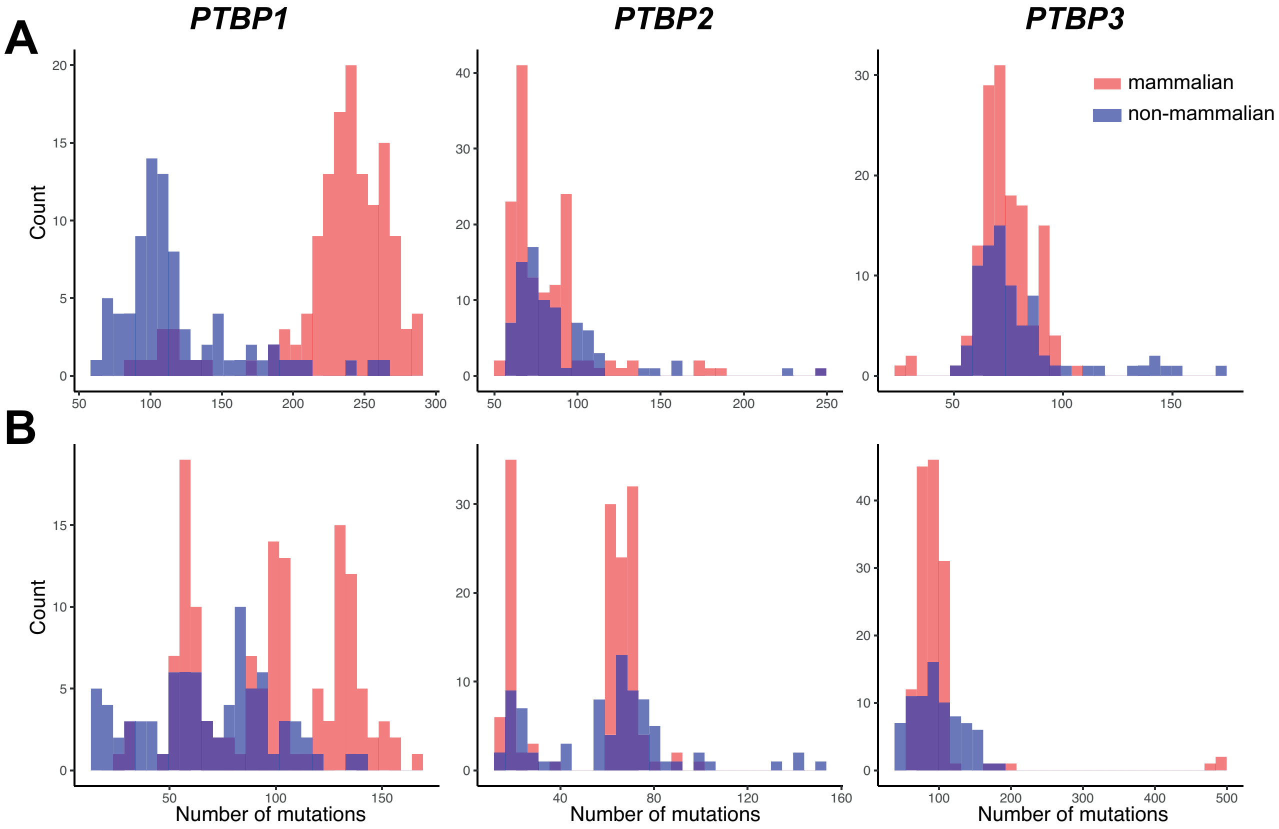


Figure S9: **Number of synonymous (A) and non-synonymous substitutions (B) for *PTBP1*, *PTBP2* and *PTBP3* mammalian (red) and non-mammalian individuals (blue).** The values presented here have been gathered through all vertebrate species studied. X-axis represents the number of substitutions between a species and its ancestral state and Y-axis represents the frequency of that occurrence.

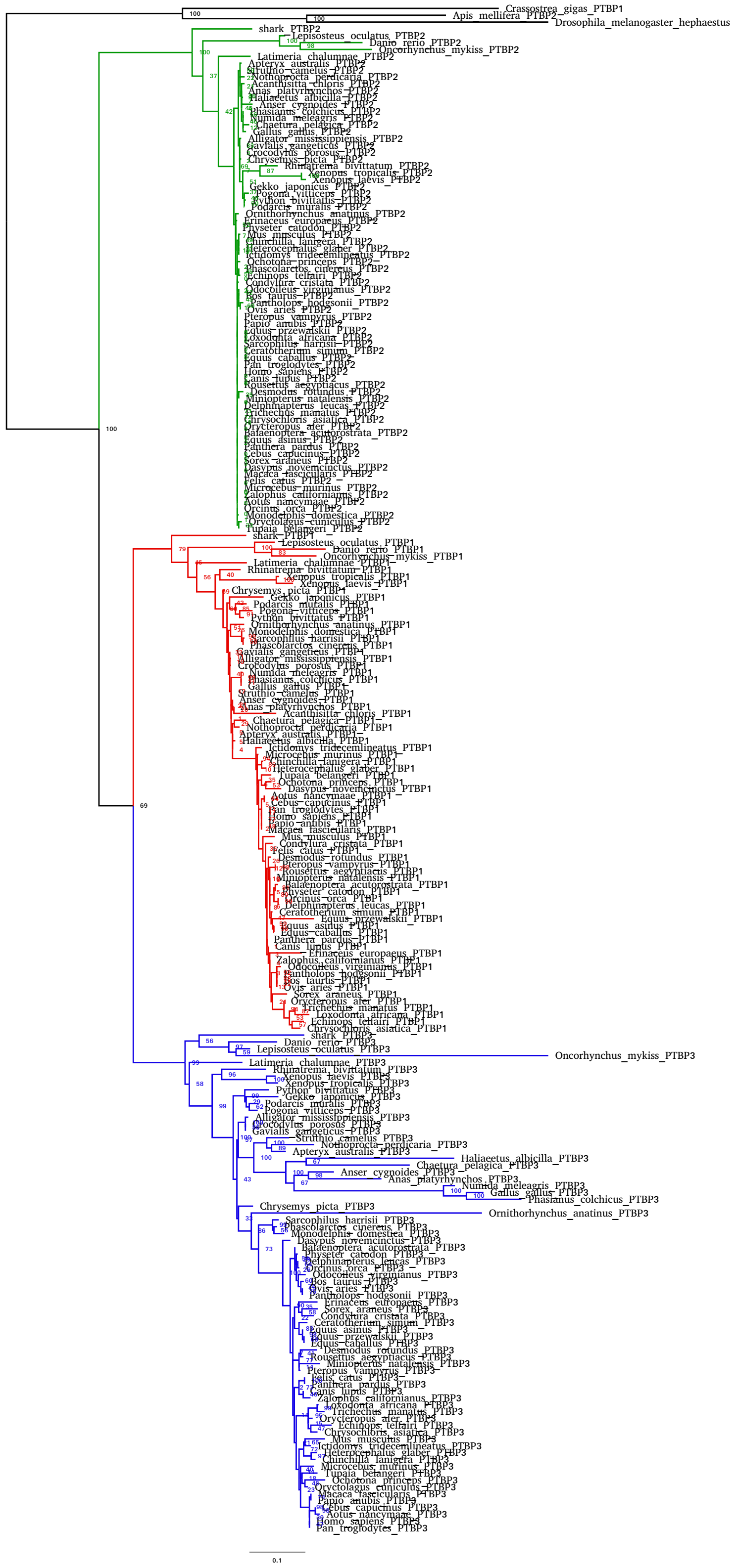

Figure S10: **Maximum-likelihood amino acid phylogeny of *PTBP*s genes.** The phylogram depicts *PTBP2*s (green), *PTBP1*s (red) and *PTBP3*s (blue) clades. The outgroup genes from protostomata are in black.

Table S11: **Synonymous and non-synonymous matrices for mammalian and non-mammalian *PTBP1*, *PTBP2* and *PTBP3*.** The matrices display cumulative counts of substitutions between each individual and their ancestral state. In each matrix, each value represents the total count of the corresponding substitution from the nucleotide in the ancestral state to the nucleotide in the extant sequences, *e.g.* there are a total of 367 A->T synonymous substitutions in *PTBP1*s in the mammalian lineages.

**Synonymous substitutions**

**Mammalians**

| ***PTBP1*** | A | T | G | C | ***PTBP2*** | A | T | G | C | ***PTBP3*** | A | T | G | C |
|---|---|---|---|---|---|---|---|---|---|---|---|---|---|---|
| A | - | 367 | 566 | 678 | A | - | 716 | 2232 | 686 | A | - | 736 | 1972 | 1186 |
| T | 334 | - | 109 | 632 | T | 774 | - | 378 | 1678 | T | 352 | - | 175 | 2380 |
| G | 3242 | 1019 | - | 631 | G | 1274 | 279 | - | 180 | G | 996 | 188 | - | 223 |
| C | 2004 | 3243 | 497 | - | C | 814 | 1465 | 67 | - | C | 538 | 1077 | 138 | - |

**Non-mammalians**

| ***PTBP1*** | A | T | G | C | ***PTBP2*** | A | T | G | C | ***PTBP3*** | A | T | G | C |
|---|---|---|---|---|---|---|---|---|---|---|---|---|---|---|
| A | - | 374 | 1188 | 626 | A | - | 401 | 1036 | 284 | A | - | 335 | 972 | 502 |
| T | 233 | - | 134 | 1140 | T | 446 | - | 155 | 836 | T | 279 | - | 78 | 1047 |
| G | 924 | 179 | - | 208 | G | 842 | 253 | - | 53 | G | 578 | 127 | - | 122 |
| C | 452 | 821 | 192 | - | C | 414 | 906 | 69 | - | C | 240 | 764 | 63 | - |

**Non synonymous substitutions**

**Mammalians**

| ***PTBP1*** | A | T | G | C | ***PTBP2*** | A | T | G | C | ***PTBP3*** | A | T | G | C |
|---|---|---|---|---|---|---|---|---|---|---|---|---|---|---|
| A | - | 108 | 814 | 336 | A | - | 96 | 478 | 94 | A | - | 327 | 1286 | 666 |
| T | 241 | - | 97 | 37 | T | 142 | - | 96 | 73 | T | 485 | - | 215 | 617 |
| G | 1098 | 270 | - | 333 | G | 406 | 51 | - | 237 | G | 1498 | 147 | - | 268 |
| C | 412 | 276 | 364 | - | C | 120 | 127 | 51 | - | C | 1170 | 411 | 196 | - |

**Non-mammalians**

| ***PTBP1*** | A | T | G | C | ***PTBP2*** | A | T | G | C | ***PTBP3*** | A | T | G | C |
|---|---|---|---|---|---|---|---|---|---|---|---|---|---|---|
| A | - | 78 | 500 | 274 | A | - | 75 | 402 | 170 | A | - | 115 | 640 | 366 |
| T | 119 | - | 80 | 76 | T | 127 | - | 109 | 49 | T | 235 | - | 130 | 386 |
| G | 302 | 67 | - | 140 | G | 346 | 48 | - | 158 | G | 818 | 86 | - | 212 |
| C | 228 | 83 | 182 | - | C | 142 | 116 | 50 | - | C | 520 | 188 | 132 | - |

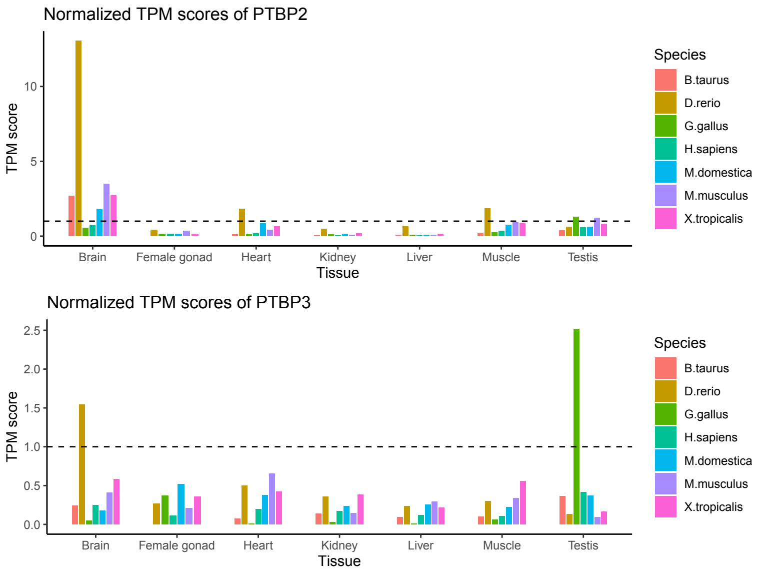


Figure S12: **Normalized expression of PTBPs in 7 species.**

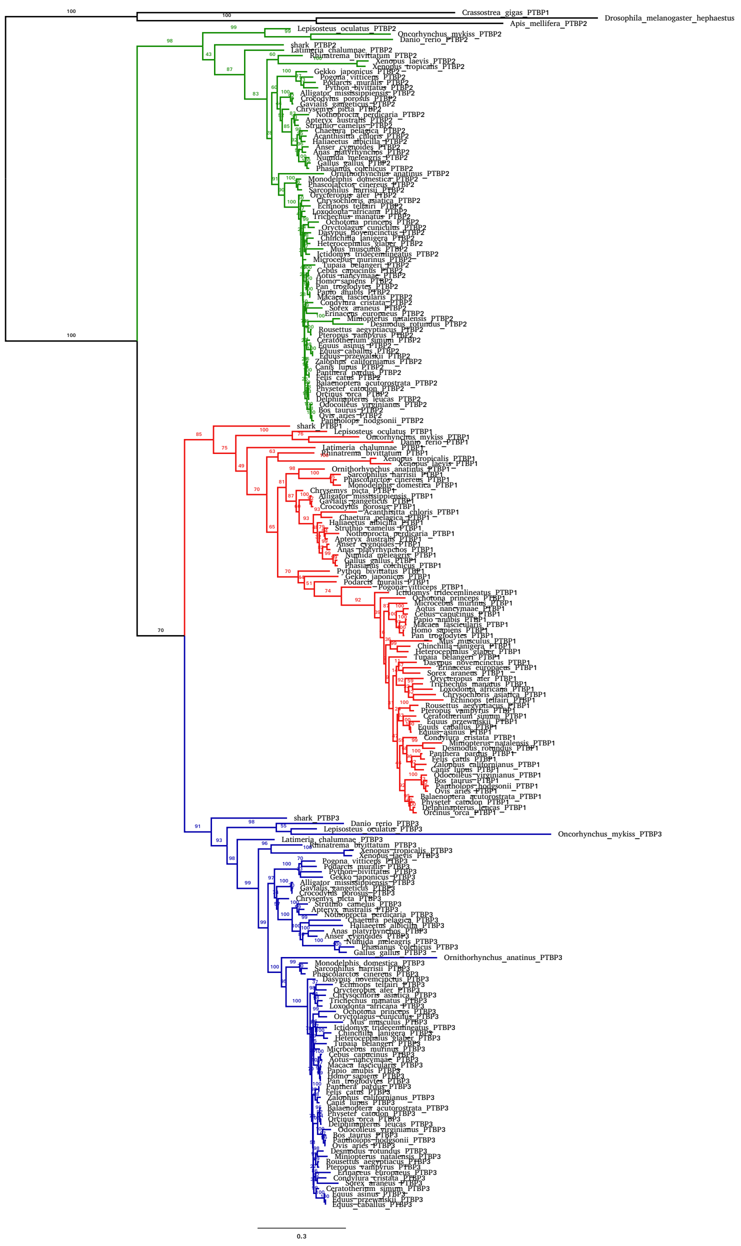


Figure S13: **Maximum-likelihood nucleotide based phylogeny of *PTBP*s genes with one-step aligement.** All sequences were aligned simultaneously. We used the GTR+GAMMA+I model for phylogenetic inference . The phylogram depicts *PTBP2*s (green), *PTBP1*s (red) and *PTBP3*s (blue) clades. The outgroup genes from protostomata are in black.

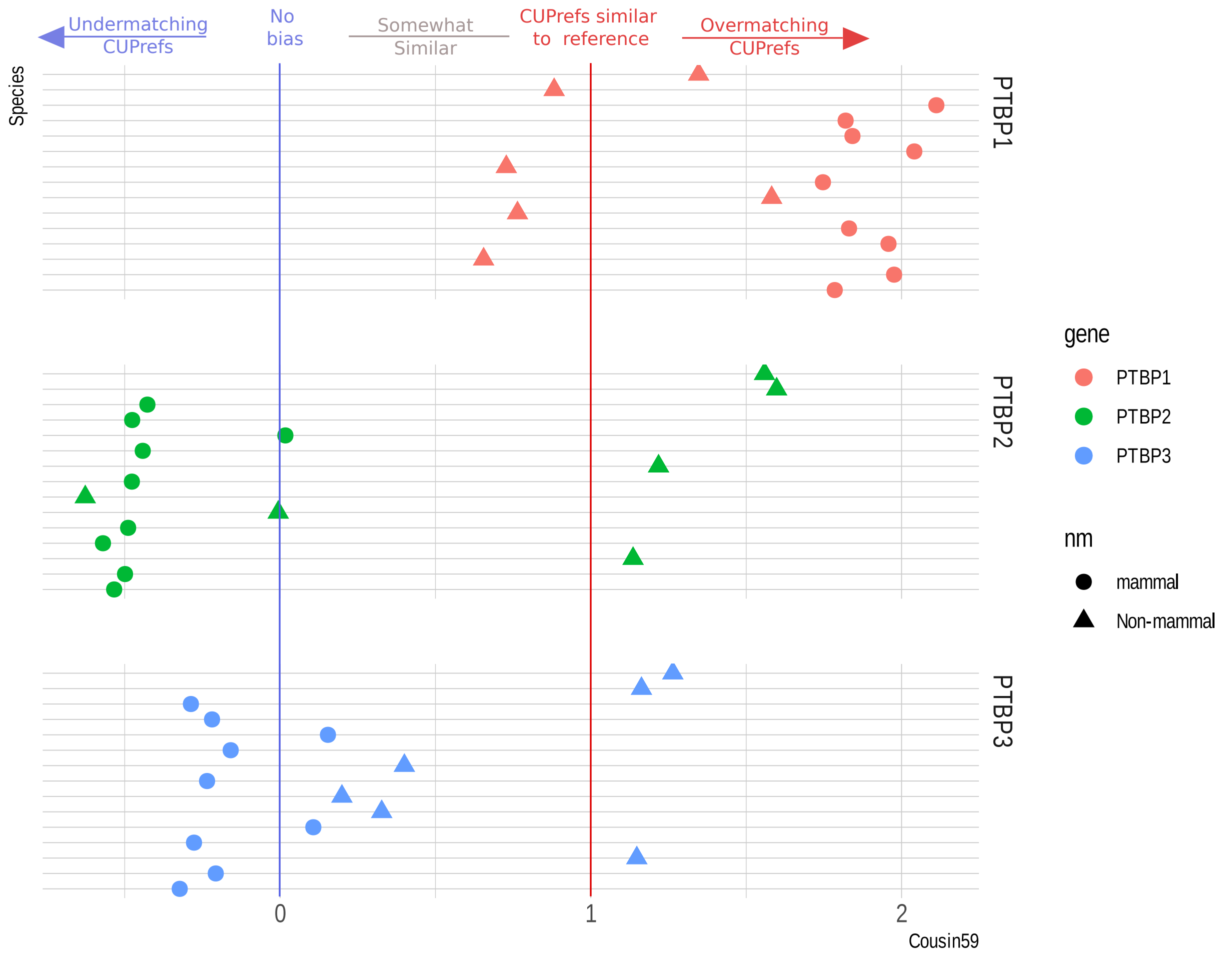


Figure S14: **Cousin59 values of the 15 selected species** by orthology (colour code for the different *PTBP*s is given in the inset) and by their taxonomy (mammals, or non-mammals). A value of 0 indicates a CUPrefs similar to a Null Hypothesis describing equal usage of synonymous codons. A value of 1 indicates a CUPrefs similar to the reference, which is here the global CUPrefs of the studied organism.

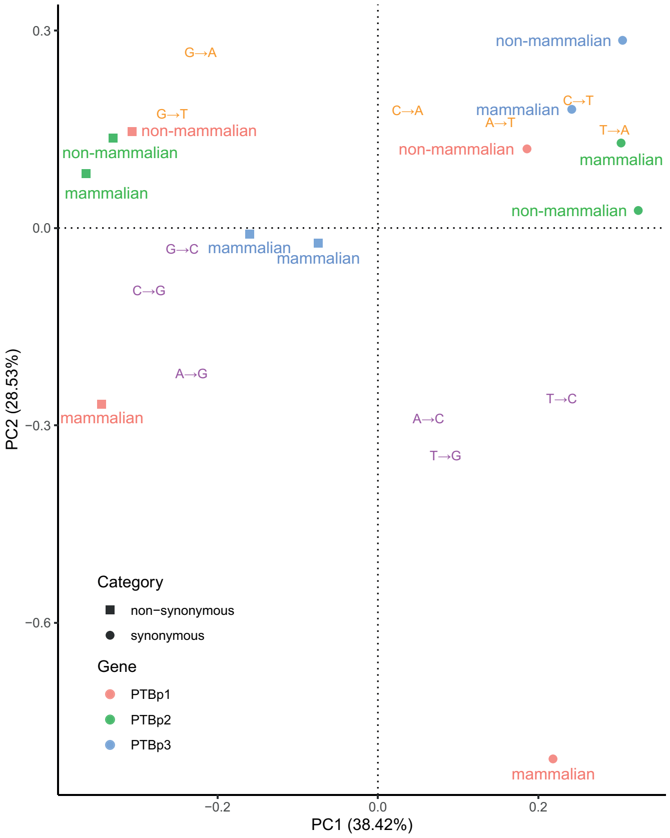


Figure S15: **Spectra of synonymous and non-synonymous substitutions for *PTBPs*.** This principal component analysis (PCA) has been built using the observed nucleotide synonymous and non-synonymous substitution matrices for each *PTBP* paralog, inferred after phylogenetic inference and comparison of extant and ancestral sequences. The variables in this PCA are the types of substitution (e.g. A->G), identified by a colour code as GC-enriching / stabilizing substitutions (purple) or AT-enriching / stabilizing substitutions (orange). Variables are plotted according to their eigenvalues. Individuals in this PCA are the substitution categories in *PTBP* genes, stratified by their nature (synonymous or non-synonymous), by orthology (colour code for the different *PTBPs* is given in the inset) and by their taxonomy (mammals, or non-mammals).

### References

Zhang Z, Li J, Cui P, Ding F, Li A, Townsend JP, Yu J. 2012. Codon Deviation Coefficient: a novel measure for estimating codon usage bias and its statistical significance. BMC Bioinformatics. 13:43.
